# Whole genome sequencing uncovers the structural and transcriptomic landscape of hexaploid wheat/*Ambylopyrum muticum* introgression lines

**DOI:** 10.1101/2021.11.16.468825

**Authors:** Benedict Coombes, John P. Fellers, Surbhi Grewal, Rachel Rusholme-Pilcher, Stella Hubbart-Edwards, Caiyun Yang, Ryan Joynson, Ian P. King, Julie King, Anthony Hall

## Abstract

Wheat is a globally vital crop, but its limited genetic variation creates a challenge for breeders aiming to maintain or accelerate agricultural improvements over time. Introducing novel genes and alleles from wheat’s wild relatives into the wheat breeding pool via introgression lines is an important component of overcoming this low variation but is constrained by poor genomic resolution and limited understanding of the genomic impact of introgression breeding programmes. By sequencing 17 hexaploid wheat/*Ambylopyrum muticum* introgression lines and the parent lines, we have precisely pinpointed the borders of introgressed segments, most of which occur within genes. We report a genome assembly and annotation of *Am. muticum* that has facilitated the identification of *Am. muticum* resistance genes commonly introgressed in lines resistant to stripe rust. Our analysis has identified an abundance of structural disruption and homoeologous pairing across the introgression lines, likely caused by the suppressed *Ph1* locus. mRNAseq analysis of six of these introgression lines revealed that novel introgressed genes are rarely expressed and those that directly replace a wheat orthologue have a tendency towards downregulation, with no discernible compensation in the expression of homoeologous copies. This study explores the genomic impact of introgression breeding and provides a schematic that can be followed to characterise introgression lines and identify segments and candidate genes underlying the phenotype. This will facilitate more effective utilisation of introgression pre-breeding material in wheat breeding programmes.

## Introduction

*Triticum aestivum* L. (bread wheat) is a vital crop, providing around 20% of calories and 25% of protein consumed globally (Reynolds et al., 2012). Improvements to wheat since the late 19th century have largely come from conventional breeding strategies, but these improvements rely on ample genetic variation in the primary gene pool (Hao et al., 2020). The hexaploid bread wheat grown today derives from just one or two polyploidisation events ^~^10,000 years ago between the tetraploid *Triticum turgidum* and the diploid *Aegilops tauschii* (Charmet, 2011). The limited diversity stemming from this genetic bottleneck has been compounded over time by intensive breeding. Pressure on breeders to prioritise advanced breeding material (J. Valkoun, 2001) for more rapid development of uniform, high quality varieties has limited the introduction of genetic variation from external sources. The genetic variation that does exist in modern wheat material is rapidly being exhausted, evident in plateauing yield improvements that left unchecked, will be insufficient to meet global demands (Ray et al., 2013). Wild relative introgression breeding will be a major component of overcoming this genetic constraint in the years to come, enabling breeders to access the secondary and tertiary gene pools of wheat (Hao et al., 2020; J. Valkoun, 2001) and incorporate novel alleles or genes into modern breeding material.

There are many examples of the successful transfer of wild relative genes into wheat since first pioneered by E.R. Sears (Doussinault et al., 1983; Fatih, 1983; Friebe et al., 1996; Klindworth et al., 2012; Sears, 1956). However, challenges associated with the high-throughput production and verification of introgression lines, in addition to the linkage drag of introgressed segments, has limited the widespread adoption of introgression breeding. Utilising recombination mutants and high-throughput marker methods, introgressing entire wild relative genomes into wheat as stably inherited, homozygous segments is now possible (King et al., 2019, 2017). These sets of lines provide the raw material required for the incorporation of alien variation into breeding programmes. Segments in these lines that confer phenotypes of interest can be identified. Lines with overlapping segments can then be crossed to break down large segments (Khazan et al., 2020), resulting in genes of interest captured in short introgressed segments with reduced linkage drag, ready to be deployed in breeding programmes.

Identifying the introgressed content of each introgression line is important for the effective utilisation of these lines. Insufficient marker density for genotyping approaches such as Kompetitive allele specific PCR (KASP) and low resolution of genomic *in situ* hybridisation (GISH) limits the resolution at which segments can be identified. Determining the precise size and positions of segments and refining positions of overlap between introgression lines is important when relating to phenotypic data to narrow down regions containing genes of interest. Identifying introgression boundaries at a higher resolution will allow lines with overlapping segments to be identified; these can be crossed to break down segments and capture genes of interest in smaller segments with reduced linkage drag.

Wild relative genes have undergone selection in a different environment to the agricultural setting in which elite wheat lines are selected and thus may be deleterious, or be, at the very least imperfect replacements of their wheat orthologue when deployed in field conditions. Therefore, many genes introgressed along with a gene of interest will contribute to reduced agronomic performance of a line. This reduced performance will be driven by differences both in the encoded protein and in the pattern of expression of the introgressed gene compared to the wheat orthologue it replaced. In addition to these direct changes to gene expression caused by introgression, disruptions to established regulatory networks and the resulting indirect effects on the expression of wheat genes in the genomic background will likely contribute to altered performance.

Ordinarily, hexaploid wheat behaves as a diploid during meiosis. The *Pairing Homoeologous 1* (*Ph1*) locus is largely responsible for this behaviour, restricting synapsis and crossovers to homologous chromosomes (Rey et al., 2017). A suppressed or deleted *Ph1* locus enables recombination between wheat chromosomes and non-homologous wild relative chromosomes and is a major tool used to transfer wild relative genes into wheat (Martín et al., 2017). However, this also enables homoeologous chromosomes to pair and recombine leading to transmission of chromatin between the subgenomes of wheat (Koo et al., 2020) and deletions/duplications where synteny between homoeologous chromosomes breaks.

Here, we have conducted a high-resolution genomic analysis on 17 *Am muticum*/hexaploid wheat introgression lines (King et al., 2019, 2017), utilising whole-genome sequencing (WGS) data from the introgression lines and the parent lines and a draft genome assembly of *Amblyopyrum muticum* [(Boiss.) Eig.; *Aegilops mutica* Boiss; 2n=2X=14; genome TT], a wild relative of wheat belonging to its tertiary gene pool. Phenotypic screening of *Am. muticum* introgression lines (Fellers et al., 2020) has revealed resistances to leaf, stem and stripe rust not observed in the parental wheat lines and thus likely conferred by introgressed genes. KASP genotyping to identify segments has been conducted on many of these lines (Grewal et al., 2021). Through this analysis, we have pinpointed introgression segment junctions to a higher resolution than previously possible, in many cases within a single pair of reads, demonstrating segments of variable size that overlap between introgression lines, which explains some differences in resistance phenotype seen between lines. These overlaps will enable these segments to be further broken down by crossing introgression lines together. Using *in silico* karyotyping, we have shown that large-scale structural disruption is ubiquitous across the lines, including deletions and duplications up to whole-chromosome size and homoeologous recombination likely facilitated by *Ph1* suppression. A genome assembly and gene annotation of *Am. muticum* has enabled us to identify introgressed resistance genes in stripe, stem and leaf rust resistant lines that may represent novel resistance conferred by *Am. muticum* genes. Analysis of gene expression of six introgression lines compared to the wheat parent lines has revealed that novel introgressed genes are less likely to be expressed than introgressed genes replacing an orthologue. Introgressed genes directly replacing a wheat orthologue show a tendency to be downregulated, with no significant balancing of the homoeologous copies in the remaining subgenomes.

## Results

### Whole genome sequencing facilitates high resolution introgression detection

To reveal *Am. muticum* segments within introgression lines using WGS data, we developed a workflow that utilises mapping coverage and single nucleotide polymorphism (SNP) information from the introgression line and the wheat parents. If a wheat segment is replaced by an *Am. muticum* segment the mapping coverage will drop in that region due to structural variation and breaks in synteny between wheat and *Am. muticum*. Due to the homozygous nature of the lines, homozygous muticum-specific SNPs are indicative of the site of introgression. Reads derived from an introgressed segment that aberrantly map to a non-introgressed region will map at the same position as the wheat reads coming from that region and result in heterozygous SNP calls with muticum-specific and wheat-specific alleles found at the same position. Therefore, to locate introgressions, we searched for genomic blocks with reduced mapping coverage, homozygous *Am. muticum*-specific SNPs and few heterozygous *Am. muticum*-specific SNPs. We identified introgressions using 1Mbp genomic windows and then defined the borders to a higher resolution using 100Kbp genomic windows. This was performed on 17 double haploid (DH) or backcrossed (BC) *Am. muticum* introgression lines from which Illumina paired-end short reads were produced to an average depth of around 5x. Figure 1A shows an example of this macro-level visualisation of introgression line DH65, which has a 51.29Mbp segment on the telomere of the short arm of chr4D, and a 139.6Mbp monosomic deletion on the short arm of chr5B. Macro-level genome plots for all lines can be seen in fig. S1.

**Fig. 1.**
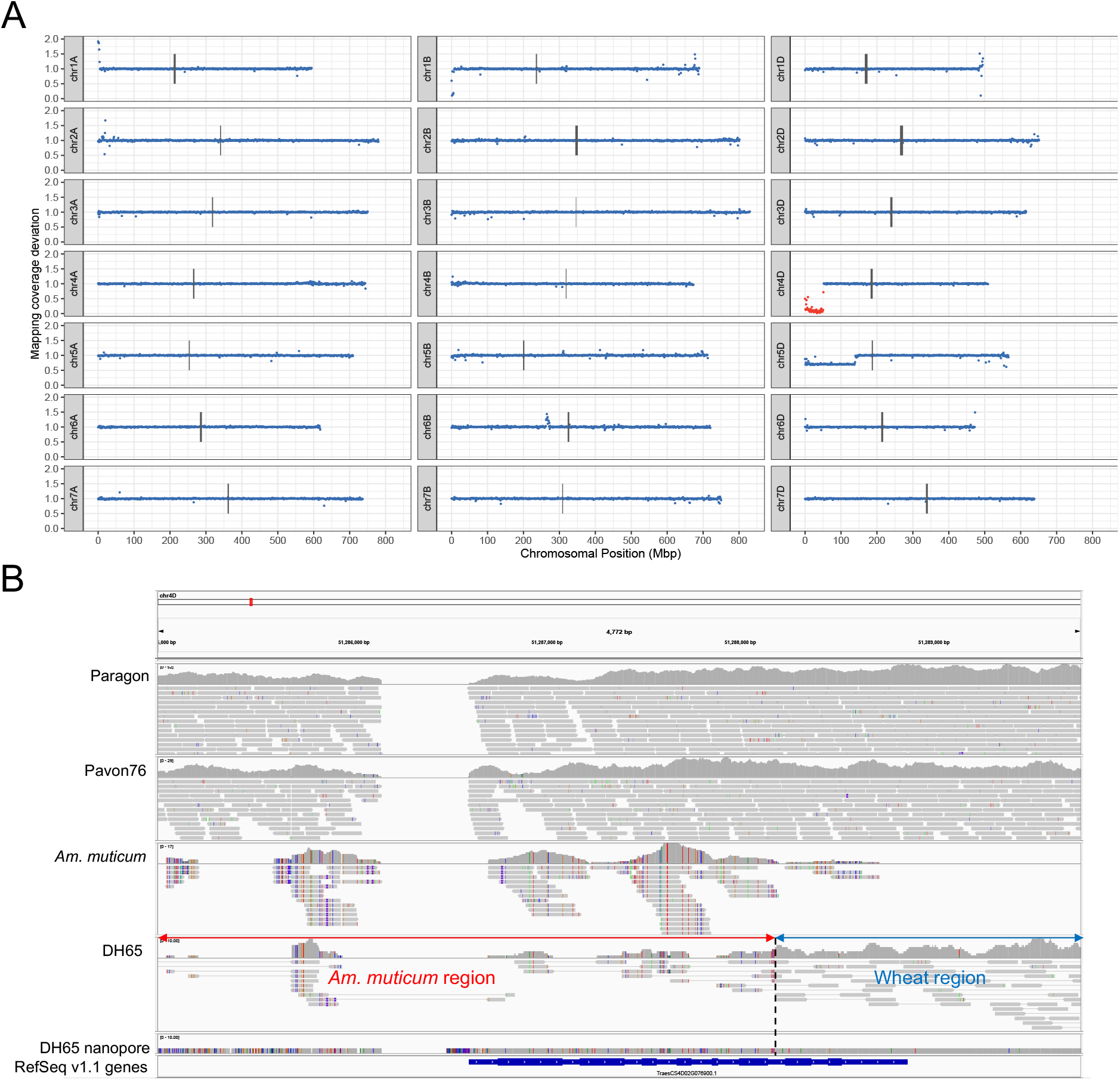
Identifying introgressed *Am. muticum* segments using whole-genome sequencing data. **A** Visualising introgressed segments and structural variation in introgression lines using WGS sequencing data. Each point represents the deviation in mapping coverage compared to the parent lines in 1Mbp windows across Chinese Spring RefSeq v1.0. Windows within assigned *Am. muticum* introgression blocks are coloured red. **A** Introgression line DH65, which has a 51.29Mbp introgressed segment on chr4D and a 139.6Mbp monosomic on chr5D. **B** IGV image showing junction at the right-hand side of chr4D segment in the introgression line DH65 (fig. 1A), spanned by both Illumina paired-end reads and Oxford Nanopore reads from DH65. The first four tracks show mapped illumina WGS data, the fifth track shows assembled contig from aligned Oxford Nanopore reads for DH65 and the bottom track shows high confidence genes from the RefSeq v1.1 annotation.

Using this approach, we confirm the existence of 100% of segments previously identified with KASP genotyping (Grewal et al., 2021). However, we were able to resolve the locations of segment junctions to a much higher resolution than previous methods, due to the limited marker density available for KASP genotyping and the inability of GISH to resolve segments below ^~^20Mbp. In addition, we were able to uncover two previously unreported segments that have been subsequently validated by KASP genotyping (Grewal et al., 2021); a 17.39Mbp on the telomere of chr7D of DH195 and a 22.68Mbp segment on the telomere of chr5D in DH121. We also identified a new 3.99Mbp segment on chr6D of DH15 that we subsequently validated using 2 KASP markers, WRC1873 and WRC1890 (Table S3). All precise segment positions are listed in Table S2.

To explore junction regions of segments in fine detail, we used the Integrative Genome Viewer (IGV) (Robinson et al., 2011), an interactive browser that allows sequencing reads mapped to a genome within a specified interval to be manually interrogated. Using IGV to explore the junction regions, we were able to precisely identify 33/42 segment ends (78.6%). As some segment ends are telomere substitutions as opposed to crossovers and some segments are derived from the same initial cross, we just looked at uniquely-derived crossover junctions and found that we could identify the precise crossover point between wheat and *Am. muticum* in 12/17 (70.6%) cases. Of the remaining junctions, two were narrowed down to within 100kbp and three had complex structures with duplication events that prohibited precise localisation. Out of the 12 high resolution junctions, 11 (91.7%) were within 670bp upstream or downstream of a wheat gene, with 8 falling within the gene body itself, suggesting that crossovers may be localised to genes. The remaining junction was 6.75Kbp downstream from the nearest gene.

For line DH65, the pinpointed junction was validated with Oxford Nanopore long reads mapped to RefSeq v1.0 along with the Illumina paired-end short reads (fig. 1B). Oxford Nanopore reads spanned the breakpoint between *Am. muticum* and wheat at the right-hand side of the 51.6Mbp chr4D segment, adding confidence to the identification from Illumina reads alone. We assembled these mapped Oxford Nanopore reads using wtdbg2 (Ruan and Li, 2020) with relaxed parameters to include reads that were clipped due to high divergence between wheat and *Am. muticum*, producing a contig that spans the junction. This contig spans the entire junction, including regions to which neither the Illumina reads from *Am. muticum* nor DH65 map. These regions appear to have elevated SNP density, explaining the gaps in mapping.

For introgression lines with KASP genotyping verification, WGS data may offer an affordable tool to aid breeders to identify precise location and size of these segments. To assess the sequencing depth requirements and locate the position and size of introgressed segments using coverage deviation alone, we downsampled the Illumina paired-end short reads from 2 lines for which we have identified very precise positions of the junction borders; DH65, and DH92, to 1x, 0.1x, 0.01x and 0.001x to choose the lowest depth at which we could still resolve segment position. 0.01x was the lowest depth that still provided comparable resolution (fig. S3).

### Introgression breeding process induces homoeologous pairing and large chromosomal aberrations

In addition to introgression sites, we have identified large deletions and duplications, many of which were whole arm or whole chromosome in scale, based on the deviations in mapping coverage not attributable to introgressions. Within the 17 lines examined, 12 lines (70.6%) have one or more very large chromosomal aberrations exceeding 140Mbp. These include: duplication of most of chr1A with a deletion of the homoeologous region of chr1B in DH pair DH124+DH355 (fig. 2A); deletion of the short arm of chr4A in DH86 and deletion of the long arm of chr4B in its DH pair, DH92 (fig. 2B); monosomic deletion of most of the short arm of chr5D in DH121 and DH65 (fig. 2C), which are not a DH pair, indicating that this event has occurred multiple times at the same position; and a monosomic deletion of chr1A in DH195 (fig. 2E).

**Fig. 2.**
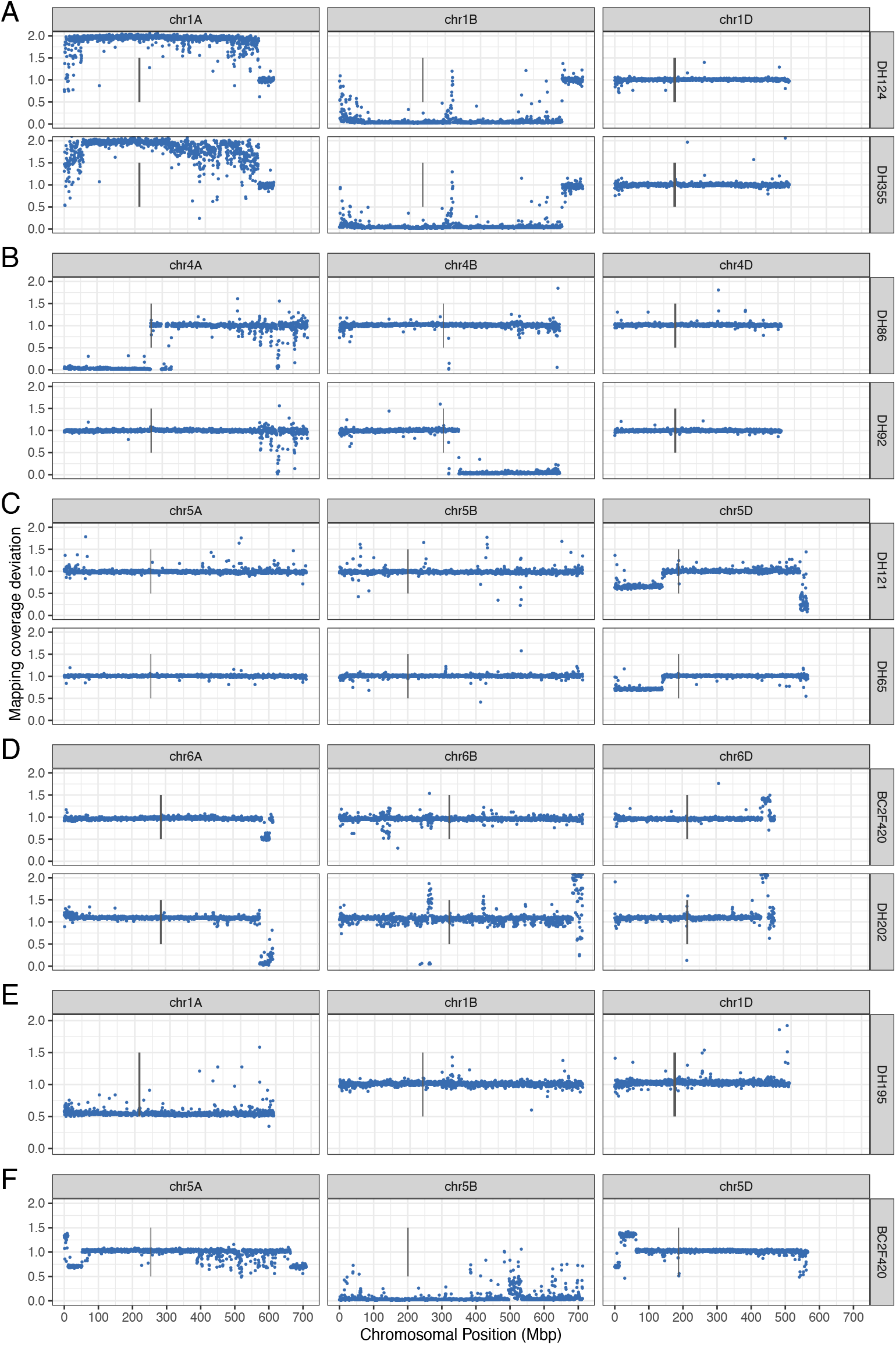
Large chromosomal aberrations in *Am. muticum* introgression lines. Each point shows mapping coverage deviation compared to the wheat parents in 500Kbp windows across the genome. **A** Corresponding duplication and deletion seen in both lines of the DH pair, caused by pairing of a duplicated chr1A and chr1B. Mapping coverage deviation of 1 at the end of chr1A and chr1B indicates a large translocation between chr1A and chr1B has taken place in duplicated chr1A+chr1B pair and discontiguous mapping coverage deviation change towards beginning of chr1A and chr1B suggests lots of smaller translocation events. **B** Chromosome arm deletions on homoeologous chromosomes of DH pair. **C** Monosomic deletions at the same position in two independently derived lines. **D** Homoeologous exchange within homoeologous group 6, at similar positions in two independently derived lines. **E** Monosomic deletion of chr1A in DH195. **F** Homoeologous recombination event between chr5A and chr5D and a deleted chr5B.

Homoeologous translocations resulting in the non-reciprocal transfer of genetic material can be detected through mapping coverage deviation, indicated by a duplication and deletion in corresponding homoeologous regions. We can also use differences within a double haploid (DH) pair (Table S1) to infer what genetic events must have taken place to give rise to the segregation patterns we see from DH lines derived from the same BC3 line. We see evidence of homoeologous pairing both from duplicated/deleted pairs of chromosomes, such as in DH355 and DH124 (fig. 2A) and from corresponding deletions/duplications at homoeologous positions (fig. 2D and fig. 2F). In BC2F420 (fig. 2F), recombination has taken place between chr5A and chr5D and chr5B has been deleted.

### Genome assembly and annotation of *Am. muticum*

To facilitate identification of introgressed genes both for differential expression analysis and to find candidate introgressed resistance genes, we produced a draft genome assembly for *Am. muticum* 2130012 comprising most of the gene space. After polishing with long and short reads and resolving haplotigs, the assembly comprised 96,256 contigs and was 2.53Gbp in length, with an N50 of 75.5Kbp (Table S5). We estimated the size of the *Am. muticum* genome through two independent methods: mapping the Oxford Nanopore reads back to the assembly and computing coverage across single-copy genes; and based on k-mer counts within the Illumina paired-end reads (fig. S4). These resulted in estimates of 4.90Gbp and 4.57Gbp, respectively, compared to flow cytometry estimate of 6.174Gbp (Pellicer and Leitch, 2020). Although the genome spans just 53.4% of the estimated genome size (mean of our two independent estimates), BUSCO analysis (Waterhouse et al., 2018) revealed that 94.2% of the expected gene space was assembled unfragmented (fig. S5). Gene annotation using evidence from root and shoot transcriptomic data, proteomic data, and ab initio predictions resulted in 86841 gene models, 32385 of which were designated as high confidence (HC) (Table 1). 28995 (89.8%) of the HC genes were assigned functional annotation.

**Table 1.** Metrics for *Am. muticum* high-confidence (HC) and low-confidence (LC) gene models

|  | HC | LC |
| --- | --- | --- |
| **Total genes (no.)** | 32385 | 54456 |
| **Single exon (no.)** | 6695 | 27364 |
| **Multi exon genes (no.)** | 25690 | 27092 |
| **Mean gene length (bp)** | 3355 | 1642 |
| **Median gene length (bp)** | 2178 | 713 |
| **Mean CDS length (bp)** | 1198 | 716 |
| **Median CDS length (bp)** | 1000 | 502 |
| **Mean exons per transcript (no.)** | 4.81 | 2.39 |
| **Median exons per transcript (no.)** | 3 | 1 |
| **Mean exon length (bp)** | 249 | 307 |
| **Median exon length (bp)** | 131 | 196 |

To identify *Am. muticum* genes not present in wheat and gene families that have undergone expansion in *Am. muticum*, both of which could be contributing novel variation in introgression lines, we used OrthoFinder (Emms and Kelly, 2019) to construct 31616 orthogroups from the proteins encoded by the HC genes from *Am. muticum, Triticum aestivum, Triticum urartu, Aegilops tauschii, Oryza sativa* and *Brachypodium distachyon* (fig. S6). 93.8% of *Am. muticum* genes were placed in an orthogroup. 3873 *Am. muticum* genes are not present in wheat and 108 orthogroups, comprising 867 *Am. muticum* genes, have undergone expansion in *Am. muticum* compared to wheat. Enrichment analysis of GO Slim terms (fig. S7) revealed that the novel *Am. muticum* genes were enriched most significantly for terms associated with metabolic processes.

### Expression of introgressed genes and impact on the background wheat transcriptome

To explore how introgressed genes are expressed and to understand the impact of the introgression breeding programme on the wheat transcriptome, we produced mRNAseq data for six of the introgression lines and the wheat parent lines. *Am. muticum* genes introgressed into each line were identified using orthologue assignments and DNA read mapping evidence. RNA reads were mapped to a pseudo genome (ABDT) constructed by concatenating the wheat reference genome, RefSeq v1.0, with the draft *Am. muticum* genome assembly; this allows us to distinguish between RNA deriving from wheat genes and from *Am. muticum* genes in the same way that we can distinguish between wheat homoeologues.

Across all six lines, 1750/4989 (35.1%) introgressed genes were expressed. Splitting the introgressed genes into those with an orthologue in wheat and those that are novel revealed that while 1627/3691 (44.1%) introgressed genes with a wheat orthologue were expressed, only 123/1298 (9.48%) novel introgressed genes were expressed (fig. 3A). For introgressed genes that do have a wheat orthologue, those that are more diverged from the orthologue are less likely to be expressed (fig. 3B), ranging from 21.5% of genes with no wheat orthologue >90% protein identity being expressed, to 64.8% of genes with an orthologue in wheat with >= 99% protein identity being expressed.

**Fig. 3.**
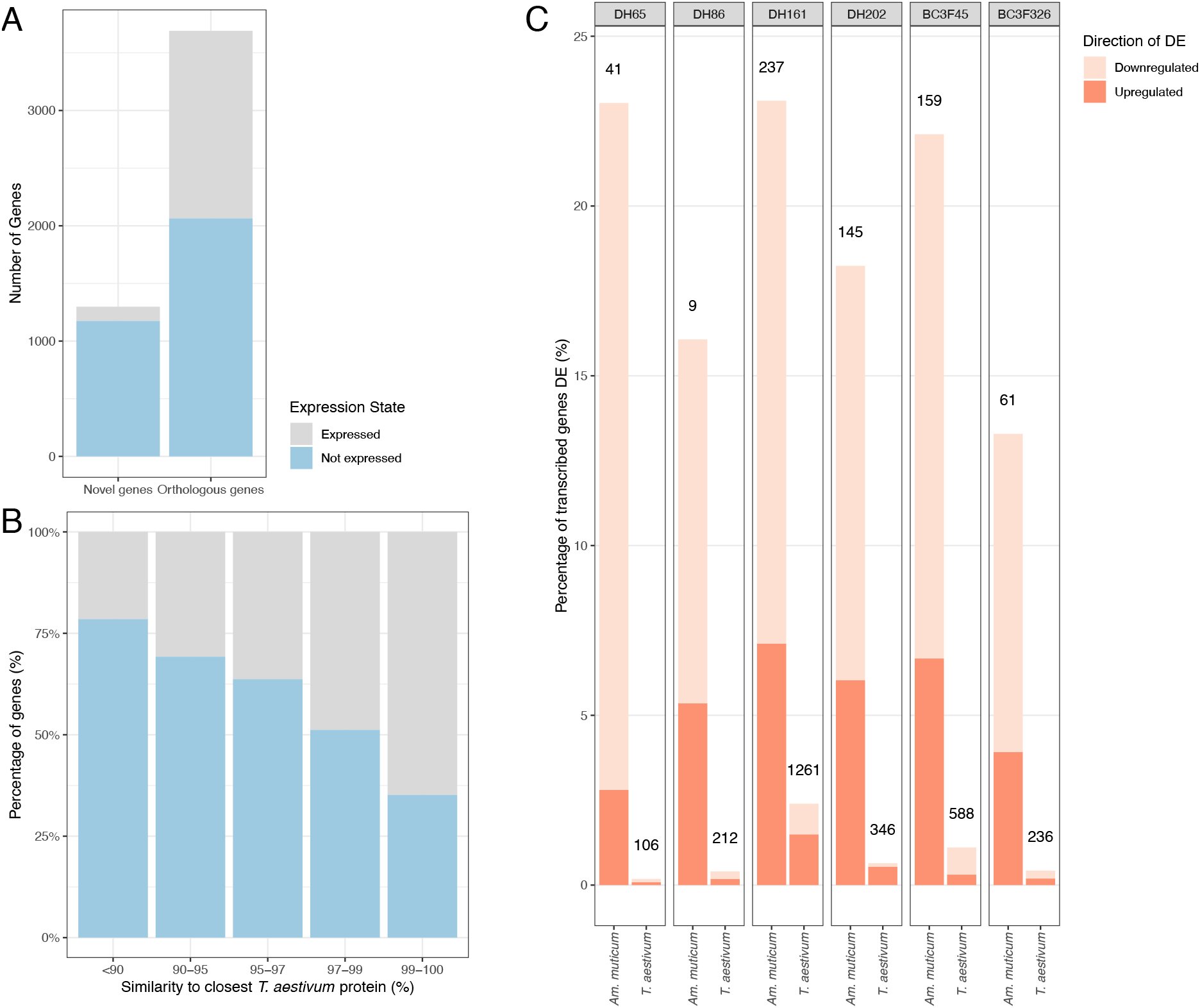
Expression of introgressed *Am. muticum* genes. **A** Expression state (Expressed or Not Expressed) of novel introgressed genes and introgressed genes in an orthogroup with a wheat gene **B** Expression state (Expressed or Not Expressed) of introgressed genes within an orthogroup with a wheat gene, binned by the protein identity between the *Am. muticum* protein and the most similar protein in the wheat reference genome annotation RefSeq v1.1. **C** Differential expression in 6 introgression lines, looking at introgressed genes compared to the orthologue they replaced in the parent lines, and background wheat genes compared to the expression in the wheat parent lines. The height of the bar represents the percentage of genes differentially expressed within introgressed and background regions for each line. The number above each bar is the number of genes called as differentially expressed.

To test whether *Am. muticum* genes that have directly replaced a wheat orthologue are expressed differently to that orthologue, we called differential expression between each introgression line and the wheat parent lines using DESeq2 (Love et al., 2014) after summing the expression count of each replaced wheat gene with that of its introgressed *Am. muticum* orthologue. Between 13.3% and 23.1% (mean of 19.3% across all lines) of introgressed genes were called as differentially expressed (abs(log_2_FC) >= 1 and adj. p-value <= 0.05 in both parental comparisons) when compared to the expression in the parent lines of the wheat orthologue they replaced (fig. 3C). Between 54.5% and 87.8% (mean of 69.8% across the lines) of these differentially expressed introgressed genes were downregulated in the introgression line.

We hypothesised that suppression of wheat genes in an introgressed or deleted region would lead to a change in the expression of homoeologous copies of that gene in the other subgenomes to compensate for the loss of expression. The results of multiple approaches support a lack of overall rebalancing of triad expression following suppression of one of the copies, both in the 400Mbp introgressed region on chr5D of BC3F45 and in the deletion of chr7D in DH161 (fig. 4). In triads where the D homoeologue has been replaced by a *Am*. muticum gene or been deleted, there is an overall reduction in expression on the D homoeologue (fig. 4A iii and 4B iii), though to a much lesser degree in the introgression where introgressed *Am. muticum* orthologues are being expressed. In the introgression, there were 74 triads with the D homoeologue introgressed and called as downregulated; none of these triads had any homoeologues called as upregulated. This was compared to 10953 control triads, where no homoeologues are introgressed or deleted and the D homoeologue is not differentially expressed, of which 17 (0.155 %) triads had the A or B homoeologue upregulated. For the deletion, out of 1294 triads with the D homoeologue deleted and therefore not expressed, just 6 triads had one or more homoeologues upregulated (0.464%); this compares to 37 (0.369%) out of 10031 control triads having the A or B homoeologue upregulated. These differences are not significant (Fisher’s exact test two-tailed p-values of 1.00 and 0.628, respectively).

**Fig. 4.**
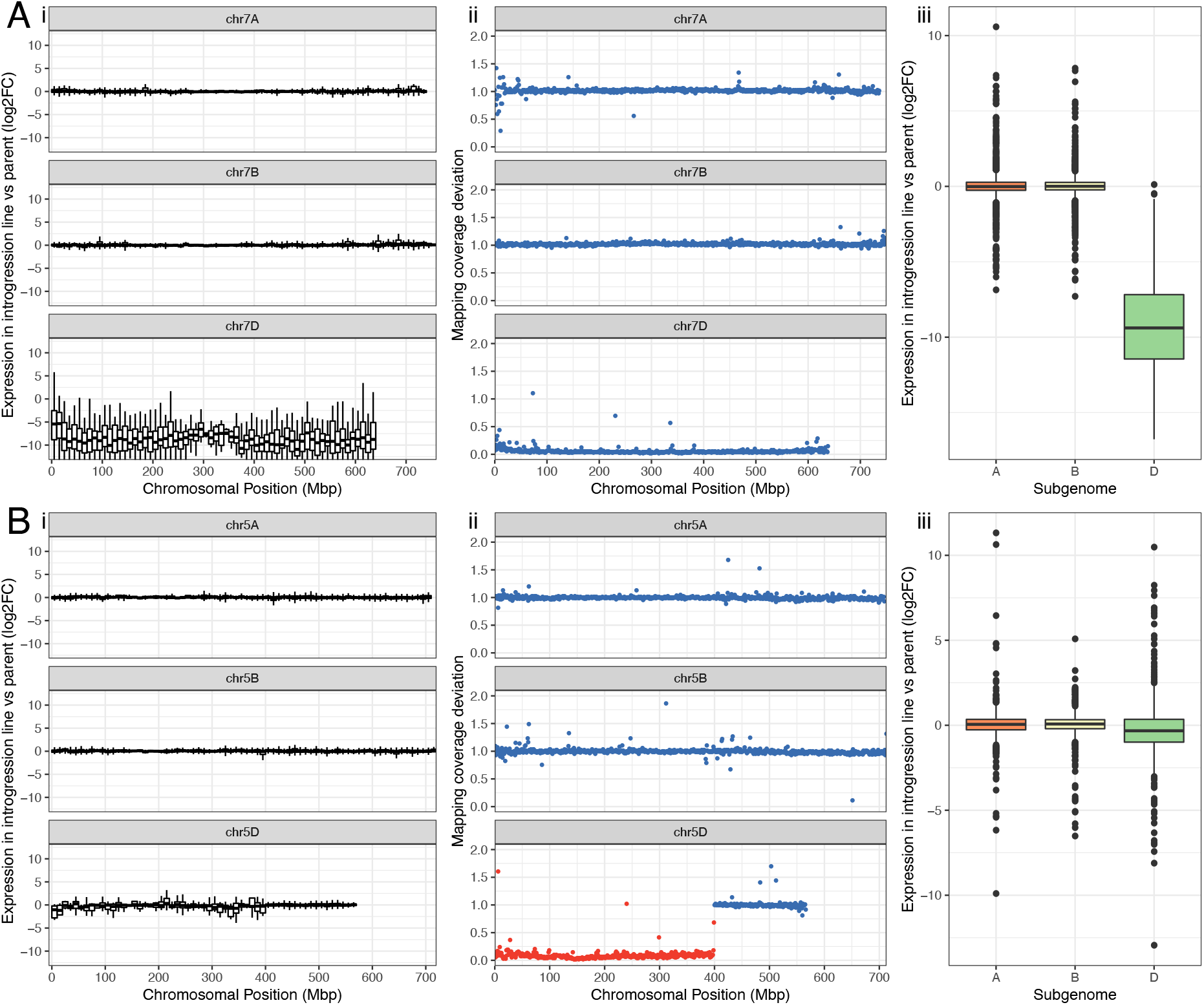
Expression profile across introgression and a deleted region and their homoeologous regions. **A** chr5A, chr5B and chr5D in BC3F45, with a chr5D:1-400Mbp introgression where chr5D genes have been replaced by *Am. muticum* orthologues **B** chr7A, chr7B and chr7D in DH161 where chr7D has been deleted. **i** DESeq2 processed log_2_FC (introgression line / Paragon) of expression compared to Paragon binned into 10Mbp window **ii** Macro level structure in 1Mbp windows. Each point represents the deviation in mapping coverage compared to the parent lines in 1Mbp windows across Chinese Spring RefSeq v1.0. Windows within assigned *Am. muticum* introgression blocks are coloured red; **iii**. log_2_FC (introgression line / Paragon) of A, B and D homoeologues belonging to triads in which the D copy has been deleted or replaced by an *Am. muticum* gene.

To complement the above approach and consider homoeologues whose expression may have changed but not sufficiently to be called as significant by DESeq2, we looked at the log_2_ fold change (log_2_FC) in DESeq2 normalised expression counts. Plotting the log_2_FC of DESeq2 normalised expression counts in 10Mbp windows (fig. 4A i and 4B i) across the chromosomes illustrates the overall stability of expression in homoeologous regions of introgressions and deletions. For the introgression and the deletion, we compared the log_2_FC of the A and B homoeologues of triads where the D homoeologue had been introgressed or deleted with the log_2_FC of the A and B homoeologues of a control set of triads defined as above (fig. S9). We found no statistically significant difference between the test and control sets (two-tailed t test p-values: deletion = 0.209; introgression = 0.252). This indicates that, like the proportion of DEGs, the change in expression counts of homoeologues in which the D homoeologue has been downregulated/silenced does not change beyond that expected by chance.

We also looked at genes in genomic windows not deviating in coverage compared to the wheat parent lines (fig. 3) to explore whether the introgressions and structural changes induced by the introgression breeding programme had indirectly affected the expression of remaining wheat genes. Between 0.181%and 2.40% (106-1261 genes; mean of 0.860% across lines) of these wheat genes were differentially expressed compared to the wheat parents. To assess whether any specific gene functions were enriched in the differentially expressed genes we looked for enriched GO Terms (fig. S8). We found some terms to be enriched, suggesting a non-stochastic impact on background transcription, however differences between lines suggests that the nature of the impact on background transcription depends on the genes introgressed/disrupted elsewhere in the genome. Some terms are enriched in more than 3 lines, suggesting these are commonly affected. These are oxidoreductase activity, oxidation-reduction process, tetrapyrrole binding, catalytic activity, carbohydrate metabolic process, cofactor binding, which are enriched in downregulated genes; and ion binding, hydrolase activity, and catalytic activity, which are enriched in upregulated background genes.

### Identifying candidate introgressed genes underlying *Am. muticum* derived rust resistance

Two of the lines that we sequenced, DH92 and DH121 (fig. 5A), have complete resistance at the seedling stage to Kansas isolates of *Puccinia striiformis tritici* (stripe/yellow rust) (Fellers et al., 2020). DH92 also displays chlorotic adult resistance to leaf rust and partial resistance to stem rust that is absent in DH121. These lines have overlapping 5D segments, the positions of which were refined to 533.2-566.1Mbp (32.9Mbp) in DH92, and 544.1-566.1Mbp (22Mbp) in DH121. Therefore, the source of the stripe rust resistance is likely within the overlapping 22.68Mbp region, and the source of leaf/stem rust resistance is likely within the 10.9Mbp region unique to DH92.

**Fig. 5.**
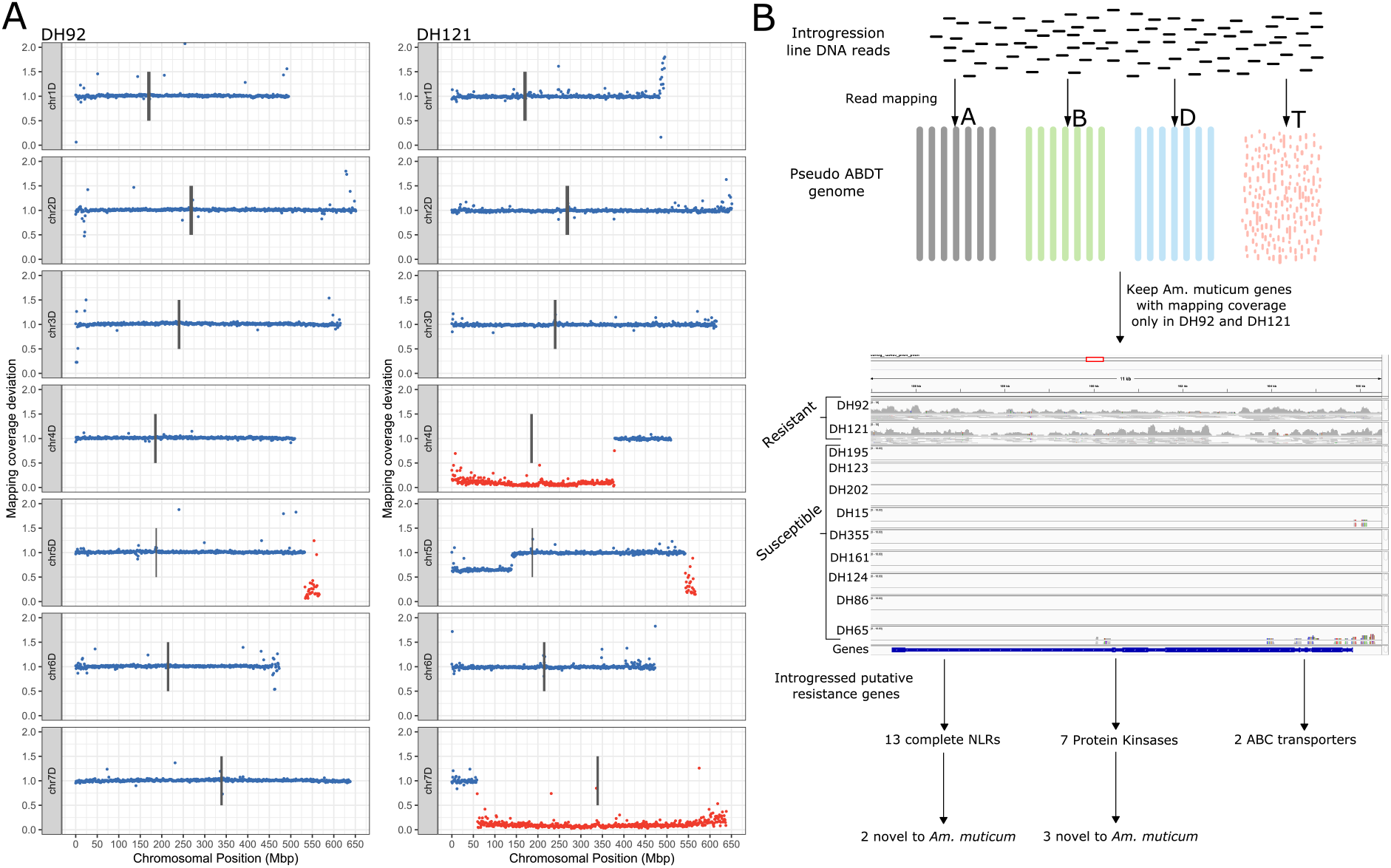
Identifying candidate introgressed resistance genes. Introgression lines DH92 and DH121 possess a partially overlapping introgressed segment on chr5D, a common resistance phenotype to stripe rust but a differential resistance phenotype to leaf and stripe rust. **A** Macro-level structure of the D subgenome of DH92 and DH121 (no segments on A or B subgenomes). Each point represents the deviation in mapping coverage compared to the parent lines in 1Mbp windows across Chinese Spring RefSeq v1.0. Windows within assigned *Am. muticum* introgression blocks are coloured red. **B** Identifying resistance genes uniquely introgressed in DH92 and DH121 and thus candidates for the stripe rust resistance shared between the two lines.

Using a mapping-based approach to the pseudo ABDT genome (fig. 5B) and combining with functional annotation, we identified 13 complete nucleotide-binding, leucine-rich repeat (NLR) immune receptors uniquely introgressed in these two lines. 12 of these have a syntenic wheat orthologue within the overlapping region of the 5D segments and 2 displayed unique NB-ARC domain signatures. 10 of the NLRs are within a 597.34Kbp cluster, including the 2 novel NLRs. We also identified 2 ABC transporters uniquely introgressed, both of which have 5D orthologues with over 97.5% protein identity, and 7 protein kinase genes uniquely introgressed, 3 of which are highly diverged at the protein level compared to the closest protein in wheat (52.2%, 74.2% and 77.0%). NLRs, ABC transporters and LRR protein kinases have all been previously implicated in resistance to stripe, leaf and stem rust (Chen et al., 2020; Krattinger et al., 2009; Wang et al., 2020). Gene candidates are detailed in Table S6.

We identified 3 wall-associated protein kinases (WAKs), and 3 protein kinases uniquely introgressed in DH92 with orthologoues or proximal to orthologues of wheat genes in the 10.9Mbp non-overlapping region of the 5D segment. 2 of the WAKs are orthologues of TaWAK388 and TaWAK390 on 5D and 1 is orthologous to TaWAK255 on chr4A. Wall-associated kinases have previously been associated with leaf rust adult plant resistance (APR) (Dmochowska-Boguta et al., 2020). Two of the protein kinases are identical at the protein level and are most similar to TaWAK387 just upstream of the TaWAK388 and TaWAK390. These may be truncated tandem duplications of this WAK. Unlike the other uniquely introgressed genes identified, the WAKs have some reads mapping to them in most of the introgression lines but only in DH92 is the coverage uniform across their lengths. This likely suggests that these are uniquely introgressed in DH92 and thus can remain as resistance candidates, but similar *Am. muticum* WAKs present in other lines are falsely mapping to these. This is supported by a lack of mapping across the rest of the contig in the other lines, unlike in DH92. Gene candidates are detailed in Table S7.

## Discussion

### Using whole-genome sequencing to pinpoint wild relative introgressions in wheat – an affordable approach to better characterise introgression lines

The current approach for studying synthetic introgression lines prior to deployment in breeding programmes relies on cytogenetic and genotyping techniques, namely GISH and KASP (Grewal et al., 2021; King et al., 2019). *De novo* discovery of SNPs to produce higher density KASP markers has improved the resolution but are insufficient for unpicking the precise size and location of segments and will likely miss small segments without the guidance of WGS data to identify areas in which additional markers should be deployed. We observe this with the new chr6D segment, the small chr7D segment in DH195 and chr5D segment in DH121, the latter two of which are sources of novel disease resistance.

We have demonstrated how whole genome sequencing data can be used to define introgressions to a very high resolution as well as resolve large-scale structural changes in these lines. Downsampling has shown that if we don’t require SNP information, only 0.01x sequencing coverage is required to pinpoint the junctions of known introgressed segments to a comparable resolution. Overlaying this information with KASP genotyping will undoubtedly provide an affordable method to characterise sets of synthetic introgression lines more accurately and comprehensively.

Introgressed segments nested within complex genomic structures, such as in DH202 (fig. S2), can only be inferred in conjunction with cytogenetic data and/or segregation patterns of DH pairs. Some introgression segment boundaries, such as the left-hand border of chr2A in DH15, can be identified but are difficult to pinpoint precisely due to structural complexities around the junction. Therefore, caution is advised when relying on introgression assignments provided by WGS data alone, particularly for complex lines with several large introgressions/deletions/duplications. However, for most lines, where genomic structure is simpler, this approach is robust, and is nevertheless an improvement on lower resolution methods, even if only to identify confounding structural complexities that would otherwise have been missed.

We have found that crossover points between wheat and *Am. muticum* mostly take place within or adjacent to genes. Previous work has shown crossovers between wheat and wild relatives are enriched in gene rich regions (Nyine et al., 2020), which mirrors recombination rates along the genome (Gardiner et al., 2019). Here we have achieved sufficient resolution to reveal these wild relative crossovers are taking place not only in regions of open chromatin and increased recombination rate, but within the genes themselves. Interestingly, this follows the same pattern previously identified for crossovers between homoeologous chromosomes (Zhang et al., 2020), in contrast to homologous crossovers which, while enriched in subtelomeric regions and at recombination hotspot motifs, are not specifically enriched in and adjacent to gene bodies.

### Genomic instability generated through introgression breeding programme

We have illustrated that structural disruption is common in introgression pre-breeding material, including homoeologous pairing and recombination, and duplications and deletions up to chromosome size. This is likely caused by the *Am. muticum*-induced suppression of the *Ph1* locus (Dover and Riley, 1972a; Dover and Riley, 1972b), however forced chromosome pairings in the F1 cross and the DH process may also be involved, although we see similar disruption in non-DH lines. An awareness of chromosomal aberrations is important for breeders using these lines in their breeding programmes. It will be important to identify the location of the *Ph1* suppressor in *Am. muticum* and other wild relatives that have an innate *Ph1* suppression system, such as *Ae. speltoides* (Li et al., 2017) to prevent segments being carried forward into breeding programmes that contain a *Ph1* suppressor that could generate further genomic disruption. For introgression lines conferring specific phenotypes of interest, it may be important to remove the chromosomal aberrations through further backcrossing, or to characterise which wheat genes have been deleted or duplicated as these may have large effects on phenotype.

Smaller scale variation in mapping coverage suggests there is structural variation taking place that we can’t accurately assess with our available data, such as transposable element mobilisation. It will be important to assess the nature and extent of such variation in the future. Unfortunately, structural variation between available chromosome-level genome assemblies and Paragon/Pavon76 is too great for structural variants arising from genome shock to be distinguished from existing structural variation between the cultivars. To study this type of variation, we will need genome assemblies of an introgression line and the wheat parents used in the cross, or a genome assembly of the wheat parent and long read data from the introgression line.

### Identification of novel introgressed genes and gene expression profile of introgression lines

We identified 3873 novel *Am. muticum* genes that could underlie novel traits to introduce into wheat. The gene expression analysis revealed that in the introgression lines, these novel genes are much less likely to be expressed than introgressed genes with an orthologue in wheat. For introgressed genes that do have an orthologue in wheat, there is a further relationship between level of divergence and likelihood of being expressed. This may reflect a lack of required regulatory elements or less efficient transcription factor binding due to divergence between the *Am. muticum* and wheat genes. However, some of this relationship could be driven by the confounding effect of more conserved genes having more core functions and therefore being more constitutively expressed (Luna and Chain, 2021). It will be important to explore this further to begin to determine whether traits identified in wild relatives may present differently when introgressed into wheat.

Many of the introgressed genes are differentially expressed compared to the wheat orthologue replaced, far exceeding the proportion of wheat genes in the background that are differentially expressed. This makes sense biologically due to the different genomic background the *Am. muticum* genes have been placed in. Two previous studies have explored the expression of genes in wheat introgression lines with barley (Rey et al., 2018) and *Aegilops longissima* (Dong et al., 2020) introgressed. Due to the differences in methods used for different studies, it is difficult to compare the total proportion of DEGs. However, both previous studies also show that many introgressed genes are differentially expressed with most of these being downregulated or silenced. Despite the elevated levels of differential expression among introgressed orthologues, it is important to note that the majority of introgressed genes replacing a direct orthologue were not differentially expressed, suggesting a remarkable similarity in expression compared to the replaced gene in the majority of cases.

We didn’t see significant change in homoeologue expression in response to introgression or deletion events. This is line with previous results showing a lack of compensation in homoeologue expression following aneuploidy (Zhang et al., 2017). This lack of response suggests that if large-scale balancing of triad expression does take place, it must require selection pressure, which these synthetic lines lack. Now that genome assemblies are available for wheat cultivars possessing many wild relatives introgressions (Walkowiak et al., 2020) that have undergone extensive artificial selection in a wheat background, it will be interesting to analyse how these introgressed regions are expressed and whether balancing of triad expression arises after a period of selection.

We see some commonly enriched GO Terms in the genomic background that may be linked with cellular stress or loss of cellular homeostasis; this conclusion is supported by conclusions drawn in the wheat/barley introgression line (Rey et al., 2018). The lines without these enriched GO Terms have less disruption overall, with fewer differentially expressed genes in the background and thus may either not have sufficient genomic stress to trigger these responses or lack sufficient sample size of DEGs to call significance.

### A case study for uncovering candidate introgressed genes underlying phenotypes of interest

Combining high resolution detection of introgressed segment borders with phenotypic information and a genic assembly of *Am. muticum* has enabled us to identify likely regions for novel rust resistances and produce lists of candidate genes. We identified the probable region of stripe rust resistance in DH92 and DH121 as being within the 22.68Mbp overlapping region of the chr5D segment. The small size and telomeric position of this segment makes it conducive for use in breeding. Within this region, we have identified candidate resistance genes, including 3 novel NLRs and 3 novel LRR Pkinase proteins. We didn’t find evidence for other classes of resistance genes that have been cloned for stripe rust resistance (Zheng et al., 2020) uniquely introgressed in these lines. The DH92 resistance to leaf rust that is not shared with DH121, is only seen in adult plants and to a composite of isolates; this race non-specific APR tends to be more durable and, in combination with the small segment size, makes this resistance another good target for further characterisation. We identified 3 WAKs and 3 protein kinases uniquely introgressed in DH92. Wall-associated kinases have previously been shown to confer resistance to leaf rust that looks similar to APR (Dmochowska-Boguta et al., 2020) and protein kinase proteins, such as Yr36, have been implicated in APR (Ellis et al., 2014). If only interested in either the stripe rust or leaf/stem rust resistance, DH92 and DH121 could be crossed to recover the desired resistance in a smaller segment with less linkage drag.

In addition to narrowing down the source of resistance genes and identifying introgressed resistance candidates, this method acts as a case study that can be built on to aid the dissection of traits in sets of introgression lines. These lines as well as many other sets of synthetic introgression lines are being phenotyped for a variety of agronomically important traits and genome assemblies for additional wild relatives are likely to be produced in the coming years. The analysis we have described here will work better with improved assemblies in which contiguous introgressed segments can be reconstructed and introgressed content fully assessed.

## Experimental procedures

### Introgression line selection

Hexaploid wheat/*Am. muticum* Introgression lines were produced as in (King et al., 2019, 2017) and summarised in method S1. 13 DH lines, 3 selfed lines and 1 heterozygous BC line, along with *Am. muticum*, Paragon, Pavon76 and Chinese Spring, were selected for DNA whole-genome sequencing (Table S1). 12 of the lines belong to a pair or a trio of lines (referred to in this manuscript as DH pairs) that derive from seed from the same BC1 cross, so common segments are not independently derived. 4 DH and 2 BC lines (Table S1), along with *Am. muticum*, Paragon, Pavon76 and Chinese Spring, were selected for RNA extraction and sequencing.

### Whole-genome sequencing, mapping and SNP calling

DNA from young leaf tissue was extracted and sequenced on Illumina NovaSeq 6000 S4 flowcells to produce 150bp paired-end reads for the introgression lines and Pavon76 and 250bp paired-end reads for *Am. muticum* (method S2). 150bp paired-end reads from Chinese Spring and Paragon was previously produced. Reads were mapped to the Chinese Spring reference genome RefSeq v1.0 (International Wheat Genome Sequencing Consortium (IWGSC) et al., 2018), followed by SNP calling and filtering (method S3).

### In silico karyotyping - calculating mapping coverage deviation compared to wheat parents

The number of mapped reads post filtering and duplicate removal was counted across genomic windows (1Mbp and 100Kbp) in RefSeq v1.0 using bedtools makewindows (Quinlan and Hall, 2010) and hts-nim-tools (Pedersen and Quinlan, 2018) for the wheat parents (Chinese Spring, Paragon and Pavon76) and each introgression line. Mapped read counts were normalised by dividing by the total read number post duplicate removal. Normalised counts of each introgression line were divided by the normalised count of each wheat parent in its crossing history (Paragon + Pavon76 or Paragon + Chinese Spring) and the number closest to 1 was kept as the coverage deviation for that window, under the assumption that the parent with mapping coverage closest to the introgression line is the parental donor in that window. The resulting number reflects the copy number of wheat DNA in that window relative to the wheat parent. A number of 1 indicates that the DNA in that window is present in the same amount as in the parent line. A number approaching 0 suggests either a deletion or an introgression has occurred at that region, and a number of 2 suggests a duplication event has taken place. Intermediate values indicate heterozygous copy number change. We defined windows with a coverage deviation between 0.8 and 1.2 as being ‘normal’ and not in copy number variation compared to the wheat parents.

### Identifying *Am. muticum*-specific SNPs and assigning introgressed regions

A set of custom python scripts were used to analyse the coverage deviation files and vcfs and identify the introgression segments in each line. These scripts, alongside more detailed methods, are available at: https://github.com/benedictcoombes/alien_detection. First, we produced 18,496,474 SNPs between *Am. muticum* and Chinese Spring that weren’t shared with either Paragon or Pavon76 (method S4). Introgression line SNPs were then assigned as *Am. muticum* if matching a muticum specific SNP in position and allele. Sites exceeding 3x mean coverage level were removed as this signifies collapsed repeat expansion. These SNPs were then split into homozygous and heterozygous and binned into 1Mbp windows using bedtools coverage (Quinlan and Hall, 2010).

Coverage deviation blocks were defined based on contiguous blocks of 1Mbp windows with coverage deviation < 0.7, with windows within 5Mbp from the previous coverage deviation block being merged. The block was discarded if < 80.0% of constituent windows had a coverage deviation < 0.7. Coverage deviation blocks were assigned as *Am. muticum* based on the presence of homozygous *Am. muticum*-specific SNPs and a high ratio of homozygous to heterozygous *Am. muticum*-specific SNPs, within 1Mbp windows across the block (method S5). Coverage deviation in 100Kbp windows either side of the larger block was used to define the borders of the segment. To locate the precise position of this junction, the BAM alignment files for *Am. muticum*, Paragon, Pavon76 and the introgression line were loaded into IGV (Robinson et al., 2011). The region around the border identified above was searched manually to find the position where the coverage and SNP profile switches from that of the wheat parents to that of *Am. muticum*.

### KASP validation

To validate the newly identified segment that hadn’t been previously validated, a KASP™ genotyping assay was conducted as described in (Grewal et al., 2020) (method S6) (Table S4).

### Junction validation using Oxford Nanopore long reads

DNA from introgression line DH65 extraction was prepared using ligation sequencing kit SQK-LSK109 and sequenced to a depth of 7x on a MinION using the R9.4.1_RevD flow cell. Reads were filtered using NanoFilt (De Coster et al., 2018) to remove reads below a quality score of 7 or a length of 1Kbp. Filtered reads were mapped to RefSeq v1.0 using minimap2 (Li, 2018) with parameters -ax map-ont and --secondary=no. Mapped reads around the breakpoint (chr4D:51283000-51595000) were extracted using samtools (Li et al., 2009), including clipped portions of mapped reads, and assembled using wtdbg2 (Ruan and Li, 2020). The resulting contigs were mapped to RefSeq v1.0 using minimap2 (Li, 2018) with parameters -ax map-ont and visualised in IGV (Robinson et al., 2011) along with the mapped Illumina paired-end short reads from the parent lines and DH65.

### Genome assembly of *Am. muticum*

DNA from *Aegilops mutica* (now *Am. muticum*) line 2130012 (JIC) was prepared using ligation sequencing kit SQK-LSK109 and sequenced on a MinION using the R9.4.1_RevD flow cell. 178Gbp of raw Oxford Nanopore long reads were filtered using NanoFilt (De Coster et al., 2018), removing reads below a quality score of 7 or a length of 1Kbp. Filtered reads were assembled using the Flye assembler (Kolmogorov et al., 2019). Following polishing integrated into Flye using Oxford Nanopore reads, we conducted 2 rounds of pilon (Walker et al., 2014) polishing using 102Gbp of Illumina paired-end short reads to correct systematic errors in the Oxford Nanopore reads. Finally, haplotigs that were not collapsed in the assembly were detected and resolved using purge_haplotigs (Roach et al., 2018). Gene completeness was assessed using BUSCO 3.0.2 (Waterhouse et al., 2018) with parameters -l viridiplantae_odb10 –species wheat and -m geno. Genome size of *Am. muticum* accession 2130012 was estimated by mapping back the Oxford Nanopore reads to putative single-copy genes and through a k-mer based approach (method S7).

### Gene annotation

Following annotation and masking of transposable elements (method S8), gene annotation was performed using *ab initio*, protein homology and transcriptome evidence from *Am. muticum* root and shoot mRNAseq data (method S9). These were sources of evidence were integrated using EvidenceModeler (Haas et al., 2008) and partitioned into high and low-confidence genes.

### Protein family analysis

OrthoFinder (Emms and Kelly, 2019) was used with default settings to cluster the longest protein encoded by high-confidence genes from *Am. muticum, Ae. tauschii, T. urartu, T. aestivum, O. sativa*, and *B. distachyon* into orthogroups. *Am. muticum* genes were classified as novel if in an orthogroup without a wheat protein or unassigned to an orthogroup. An orthogroup was determined to have expanded in *Am. muticum* compared to wheat if the orthogroup contained 4 or more *Am. muticum* proteins more than twice the number of proteins than wheat.

### Assigning orthologue pairs

First, we computed best reciprocal blast hits between *Am. muticum* and each wheat subgenome independently. *Am. muticum* proteins (extracted and translated from gff) and wheat proteins (taken from IWGSC 1.1 pep.fa file) were aligned reciprocally using blastp (Camacho et al., 2009) with parameters -outfmt 6 -max_hsps 3 -max_target_seqs 3 -evalue 1e-6. Hits were retained if percentage identity >= 90.0% and alignment length was >= 80.0% query length. *An Am. muticum* gene was placed in an orthologue pair with a wheat gene if it was in an orthogroup with that gene and the pair were each other’s best reciprocal blast hit.

### Classifying introgressed genes

The wheat reference genome RefSeq v1.0 and the draft *Am. muticum* assembly were concatenated to form a pseudo ABDT genome. Illumina paired-end short reads from the introgression lines were mapped to this genome and filtered using the same process as mapping to RefSeq v1.0 alone. Introgressed *Am. muticum* genes in each line were defined as those with mean depth across their length >= 13.2x in DH202 and >= 3x for the remaining lines (>= ^~^0.6 * mean sequencing depth) from the ABDT pseudo genome mapping above and on a contig/scaffold with a gene assigned to an orthologue pair with a wheat gene whose start position is within a region labelled as a *Am. muticum* introgression and also passes the coverage threshold above. This is a conservative classification to prevent inclusion of non-introgressed genes.

### mRNA extraction, sequencing, alignment, and quantification

mRNA was extracted and sequenced in triplicate from leaf tissue of six introgression lines, CS, Paragon, Pavon76 (method S10). RNA reads were trimmed using Trimmomatic (Bolger et al., 2014) with the parameters ILLUMINACLIP:BBDUK_adaptor.fa:2:30:12 SLIDINGWINDOW:4:20 MINLEN:20 AVGQUAL:20. The gff3 for the high confidence CS genes was concatenated with the gff3 for *Am. muticum* genes. Splice site hints for HISAT2 were produced using extract_splice_sites.py from HISAT-2.0.4 (Kim et al., 2019). The trimmed reads were mapped to the pseudo ABDT genome using HISAT2 with the splice hint file provided and parameters -k 101 --dta --rna-strandness RF. Non uniquely mapping reads were removed using samtools view -q 40. Stringtie (Pertea et al., 2015) was used to compute gene-level abundances, outputting both raw counts and transcript-per-million (TPM) values.

### Expression of introgressed *Am. muticum* genes

The protein sequences encoded by introgressed *Am. muticum* genes were aligned to the proteins encoded by RefSeq v1.1 HC genes using blastp (Camacho et al., 2009). The identity of the best hit for each protein was retained, with an identity of 0 assigned to proteins with no hit. TPM values for each gene were taken as the mean of the three replicates. Genes with mean TPM greater than 1.0 were classified as expressed.

### Differential expression analysis

For each wheat gene in a region identified as introgressed, they were either removed if not in an orthologue pair with an introgressed *Am. muticum* gene or their expression count was summed with that of its *Am. muticum* orthologue. Differential expression analysis between each introgression line and its two wheat parents was performed using DESeq2 (Love et al., 2014). A gene was classified as differentially expressed if it had an adjusted p-value below 0.05 and an absolute log_2_FC >= 1 in both parental comparisons. Differentially expressed genes were partitioned into those in introgressed regions, and in the unaffected wheat background where coverage deviation is between 0.8 and 1.2.

### Testing triad expression balancing

To examine whether genes belonging to triads that have homoeologues that have been replaced by a *Am. muticum* gene or have been deleted, we took a test set of triads (Ramírez-González et al., 2018) that satisfied the following conditions: the D copy is introgressed or deleted and called as downregulated; the A and B homoeologues are in normal copy number regions (coverage deviation between 0.8 and 1.2); and all homoeologues have normalised expression count across samples >= 1. These were compared to control sets of triads that satisfied the same conditions except the D homoeologue was within a normal copy number region and was not called as differentially expressed. These sets were used for both the comparison of number of triads with A and/or B homoeologue upregulated and for the comparisons of the mean log_2_FC of the A and B homoeologues between the test and control sample of triads. The significance of these comparisons was tested using two-tailed Fisher’s exact test and two-tailed t test, respectively.

### GO Term analysis

We transferred functional GO Term annotation from genes in the RefSeq v1.0 annotation to genes in the RefSeq v1.1 annotation if they shared greater that 99% similarity across greater than 90.0% of their length. Statistically enriched GO Terms within the differentially expressed background gene set were computed using the R package topGO (Alexa and Rahnenfuhrer, 2020) with the following parameters: nodeSize=10; classicFisher test p < 0.05 and algorithm=”parentchild”. Enrichment for GO Terms involved in biological processes was tested against all background genes that fall within windows with mapping coverage deviation between 0.8 and 1.2. For novel *Am. muticum* genes, GO terms were extracted from the eggnog functional annotation and converted to GO Slim terms using owltools Map2Slim (https://github.com/owlcollab/owltools). Enrichment was performed as above but against all *Am. muticum* HC genes.

### Identifying introgressed resistance genes

Potential resistance genes in the *Am. muticum* assembly, including NLRs, Protein Kinases and ABC transporters were identified (method S11). Resistance genes were manually checked using IGV to identify candidates with even sequencing coverage across the genes in DH92 and DH121 only, in the case of the shared stripe rust resistance, and across the genes in DH92 only, in the case of the DH92-specific leaf and stem rust resistance. To reduce the number of genes to manually check, we removed any genes with less than 2x mean mapping coverage across their length in either DH92 or DH121. The gene models were manually curated using the available evidence. For NLRs revealed by NLRAnnotator (Steuernagel et al., 2020) with no gene model but transcriptomic and *ab initio* evidence, gene models were manually constructed. The novelty of the uniquely introgressed NLRs was tested by extracting the NB-ARC domains using hmmscan (Finn et al., 2011) and aligning them using blastp (Camacho et al., 2009) to the proteins of HC genes from 10 wheat cultivars (Walkowiak et al., 2020). Hits below 85% identity were considered novel. The novelty of the other protein types was tested by aligning the whole amino acid sequence to the same protein set; here, hits below <80.0% were considered novel.

## Data availability

Sequencing data produced as part of this study, along with the *Am. muticum* assembly is available at: https://opendata.earlham.ac.uk/wheat/under_license/toronto/Hall_2021-10-08_wheatxmuticum. *Am. muticum* Illumina short-read sequencing reads available at: https://opendata.earlham.ac.uk/wheat/under_license/toronto/Grewal_et_al_2021-09-13_Amybylopyrum_muticum/. The Chinese Spring sequencing data used is available from ENA (study PRJNA393343; runs SRR5893651 and SRR5893652). The Paragon sequencing data used is available from ENA (study PRJEB35709; runs ERR3728451, ERR3760033, ERR3760405 and ERR3728448). Custom scripts used for introgression detection are available at: https://github.com/benedictcoombes/alien_detection.

## Acknowledgements

We would like to acknowledge BBS/E/T/000PR9816 (NC1 - Supporting EI’s ISPs and the UK Community with Genomics and Single Cell Analysis) for data generation and BB/CCG1720/1 for the physical HPC infrastructure and data centre delivered via the NBI Computing infrastructure for Science (CiS) group.

## Funding

BBSRC Core Capability Grant BB/CCG1720/1 (AH, RJ, RR-P)

BBSRC funded Norwich Research Park Biosciences Doctoral Training Partnership grant BB/M011216/1 (BC)

BBSRC Designing Future Wheat grant BB/P016855/1 and its constituent work packages DFW WP4 Data Access and Analysis (AH, RJ, RR-P, JK, IPK, SG, SE, CY)

BBSRC grant BB/J004596/1 as part of the Wheat Improvement Strategic Programme (WISP) (JK, IPK, SG, SE, CY)

USDA-ARS CRIS 3020-21000-011-000-D (JF)

## Author contributions

Conceptualization: AH, JL, IPK, BC

Methodology: BC, RJ, RR-P

Formal analysis: BC

Investigation: JF, SG, CY, SE

Writing - Original Draft: BC

Writing - Review & Editing: BC, AH, RR-P, JK, JF

Visualization: BC

Supervision: AH

Funding acquisition: AH, JK, IPK

## Supporting information

### Supporting figures

Figure S1: Whole genome macro-level plot for all 17 hexaploid wheat/*Am. muticum* introgression lines.

Figure S2: Macro structure of chr7A, chr7B and chr7D in four introgression lines (two DH pairs: DH195+DH202, and DH121+DH123).

Figure S3: Minimum required sequencing depth to uncover introgressed segments in introgression lines.

Figure S4: k-mer distribution of *Am. muticum* Illumina paired-end short reads used to estimate genome size.

Figure S5: BUSCO results using the viridiplantae_odb10 dataset after each sequential round of the assembly.

Figure S6: Interspecies intersection of orthogroups produced by OrthoFinder.

Figure S7: GO Slim terms enriched in novel *Am. muticum* genes.

Figure S8: GO terms enriched in differentially expressed wheat genes.

Figure S9: Testing compensation in homoeologue expression following deletion or introgression.

### Supporting tables

Table S1: Introgression lines sequenced in this study.

Table S2: Segments identified in each introgression line included in this study. If junction within or nearby a gene, the gene name is included.

Table S3: Genotyping results of DH15 with newly discovered small 6D segment. ‘a’ indicates presence of homozygous wheat-specific alleles; ‘b’ indicates presence of homozygous *Am. muticum*-specific alleles; ‘-’ indicates absence of a genome-specific allele for a particular KASP assay.

Table S4: Primer details for the KASP assays used for genotyping of DH15.

Table S5. Metrics of *Am. muticum* genome assembly.

Table S6. Potential resistance genes uniquely introgressed in DH92 and DH121, which share resistance to stripe rust.

Table S7. Potential resistance genes introgressed in DH92 but absent from DH121 and the other lines. DH92 has stem and leaf rust resistance not seen in DH121.

### Supporting experimental procedures

Methods S1: Introgression line production

Method S2. DNA extraction and whole-genome sequencing

Method S3. Read mapping and SNP calling

Method S4. Producing *Am. muticum*-specific SNPs

Method S5. Assigning coverage deviation blocks as *Am. muticum*

Method S6. KASP genotyping

Method S7. Estimating genome size

Method S8. Repeat annotation and masking

Method S9. Gene annotation

Method S10. mRNA extraction and sequencing

Method S11. Identifying resistance genes

