## Supplementary figures for "Whole genome sequencing uncovers the structural and transcriptomic landscape of hexaploid wheat/*Ambylopyrum muticum* introgression lines"

**Fig. S1.**

DH8

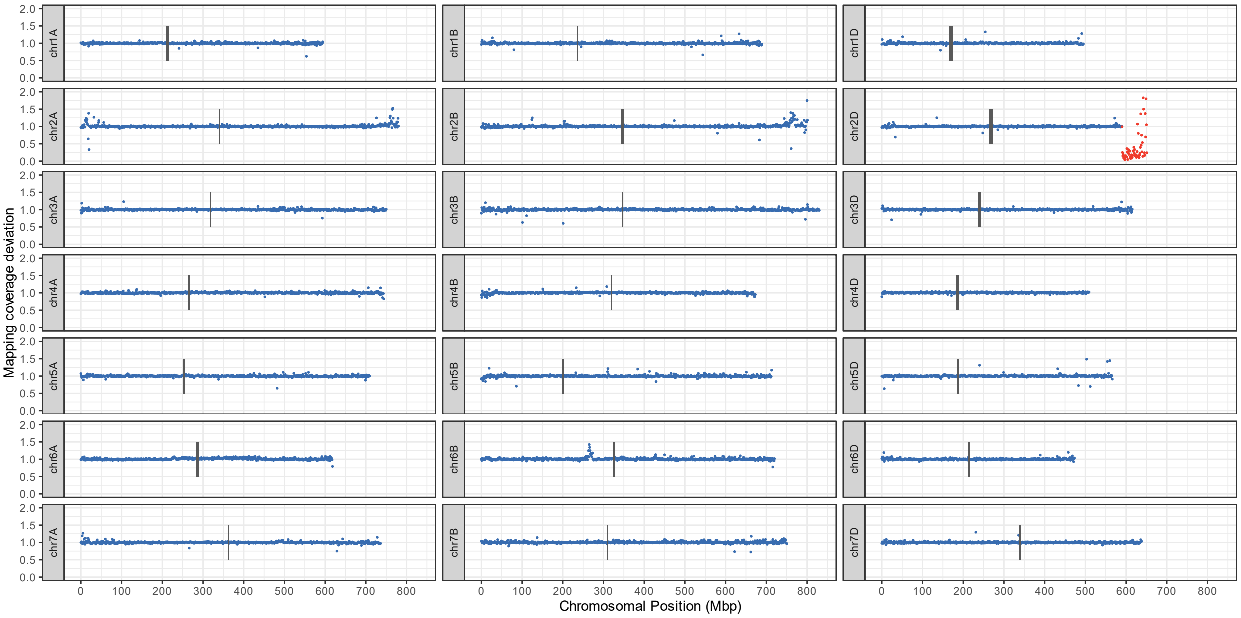


DH15

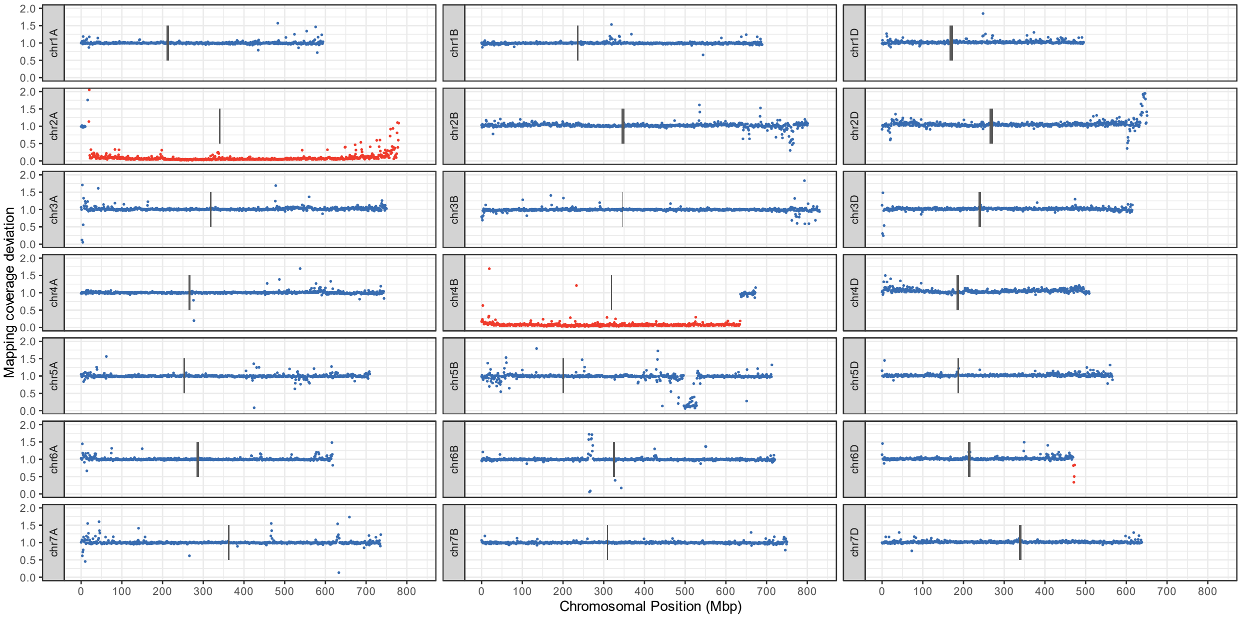

DH65

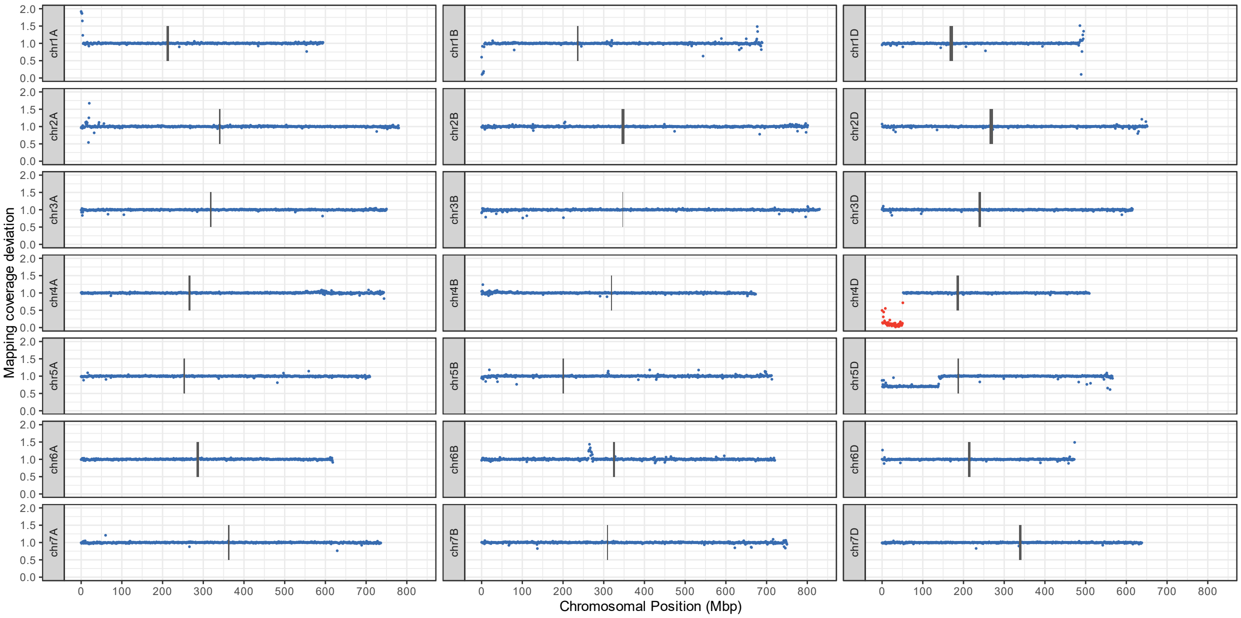
Mapping coverage deviation
Chromosomal Position (Mbp)
chr1A
chr1B
chr1D
chr2A
chr2B
chr2D
chr3A
chr3B
chr3D
chr4A
chr4B
chr4D
chr5A
chr5B
chr5D
chr6A
chr6B
chr6D
chr7A
chr7B
chr7D

DH86

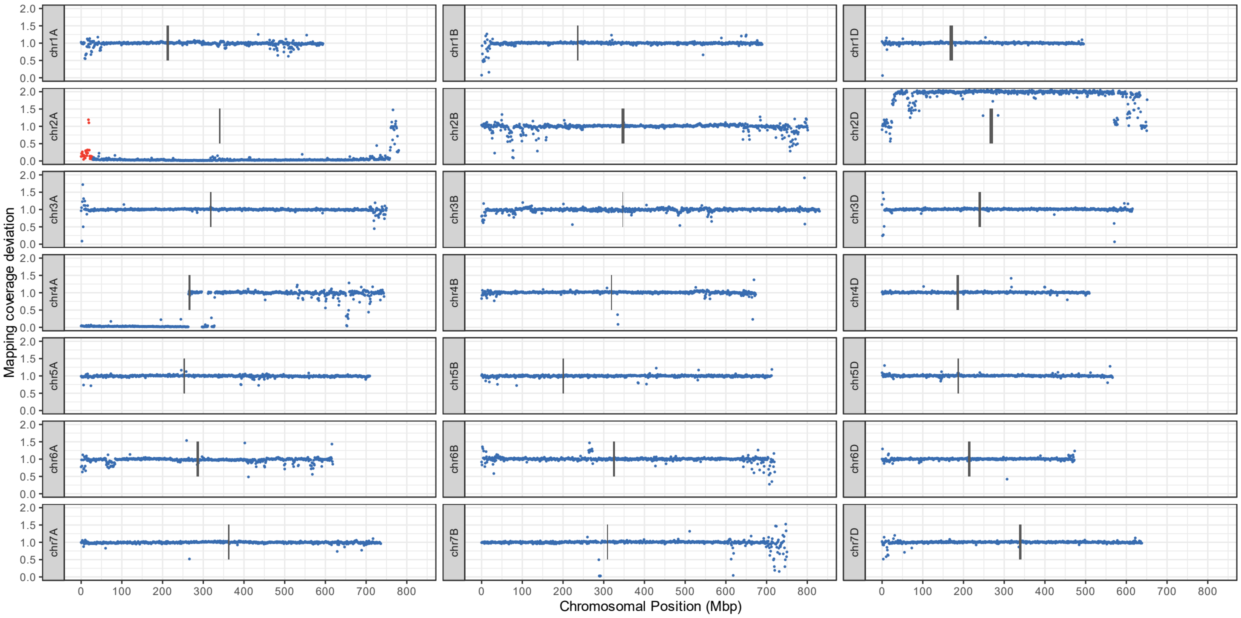
Mapping coverage deviation
Chromosomal Position (Mbp)
chr1A
chr1B
chr1D
chr2A
chr2B
chr2D
chr3A
chr3B
chr3D
chr4A
chr4B
chr4D
chr5A
chr5B
chr5D
chr6A
chr6B
chr6D
chr7A
chr7B
chr7D

DH92

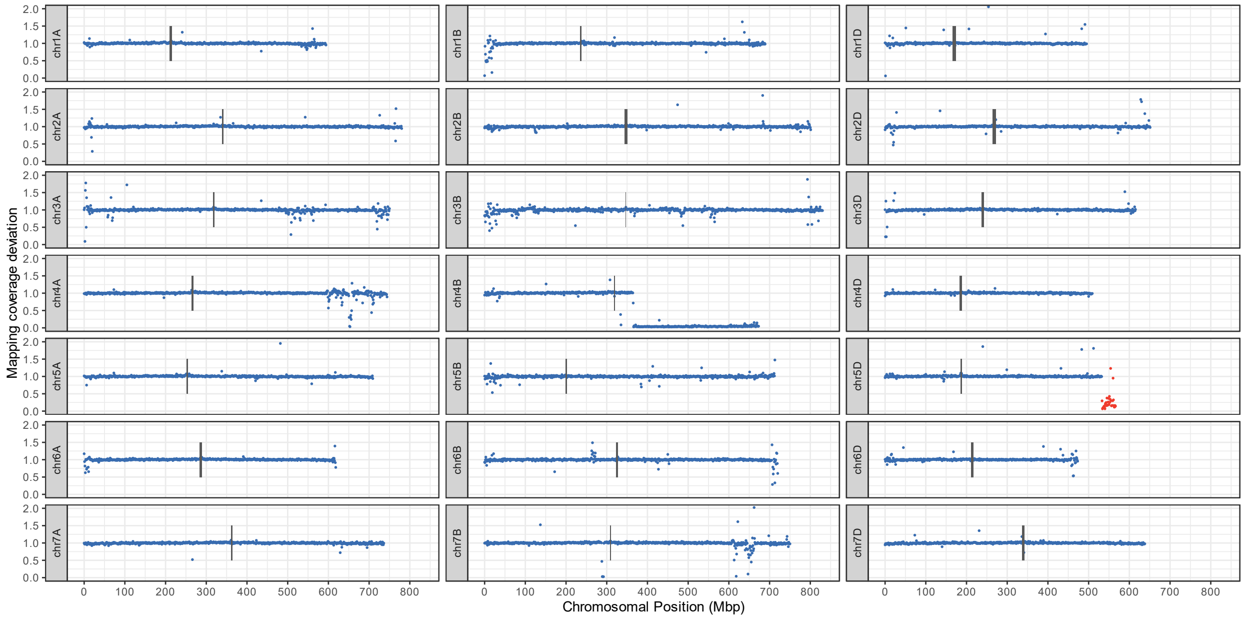
Mapping coverage deviation
Chromosomal Position (Mbp)
chr1A
chr1B
chr1D
chr2A
chr2B
chr2D
chr3A
chr3B
chr3D
chr4A
chr4B
chr4D
chr5A
chr5B
chr5D
chr6A
chr6B
chr6D
chr7A
chr7B
chr7D
0.0 0.5 1.0 1.5 2.0
0 100 200 300 400 500 600 700 800

DH96

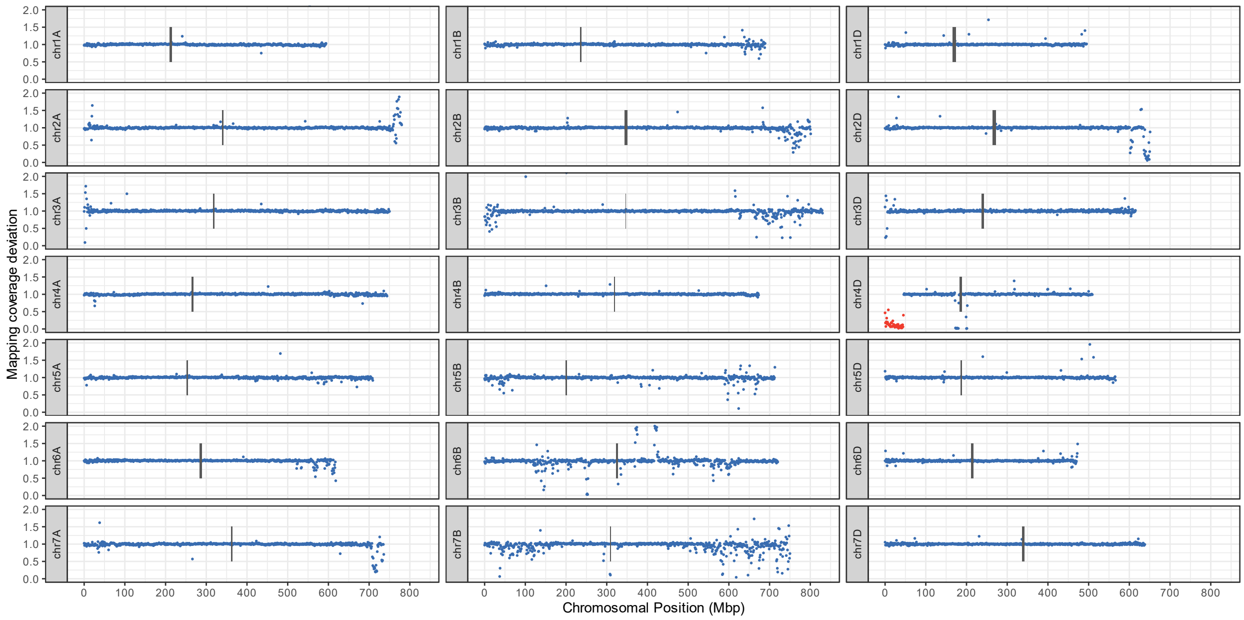
Mapping coverage deviation
Chromosomal Position (Mbp)
chr1A
chr1B
chr1D
chr2A
chr2B
chr2D
chr3A
chr3B
chr3D
chr4A
chr4B
chr4D
chr5A
chr5B
chr5D
chr6A
chr6B
chr6D
chr7A
chr7B
chr7D
0.0 0.5 1.0 1.5 2.0
0 100 200 300 400 500 600 700 800

DH121

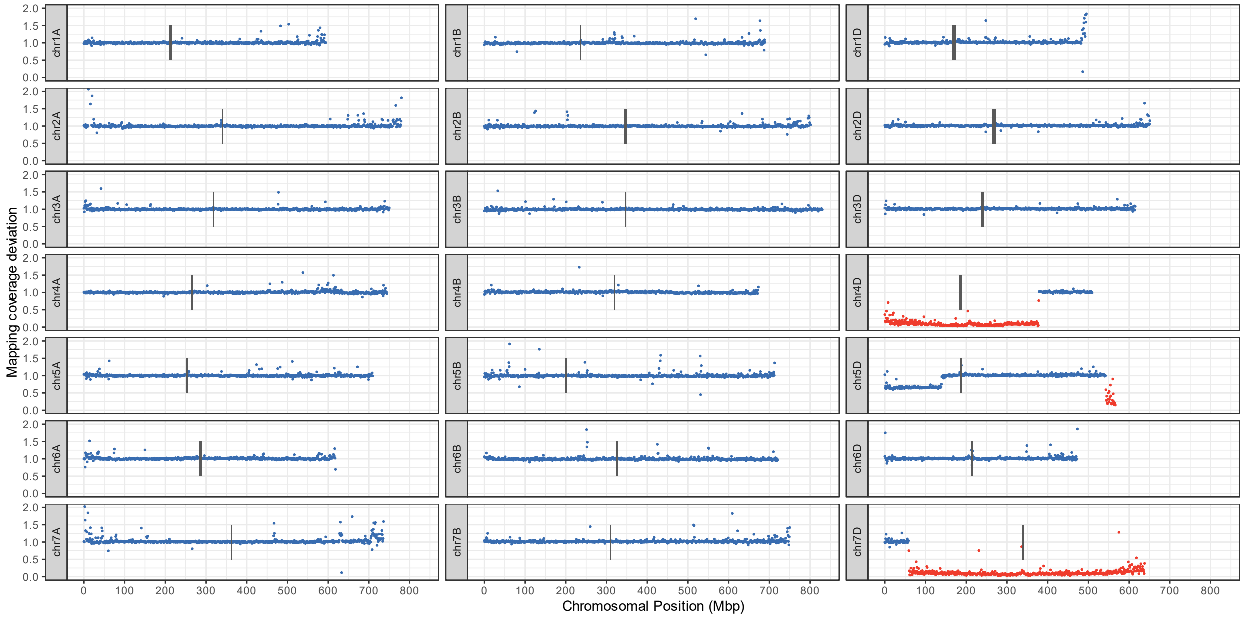
Mapping coverage deviation
Chromosomal Position (Mbp)
chr1A
chr1B
chr1D
chr2A
chr2B
chr2D
chr3A
chr3B
chr3D
chr4A
chr4B
chr4D
chr5A
chr5B
chr5D
chr6A
chr6B
chr6D
chr7A
chr7B
chr7D

DH123

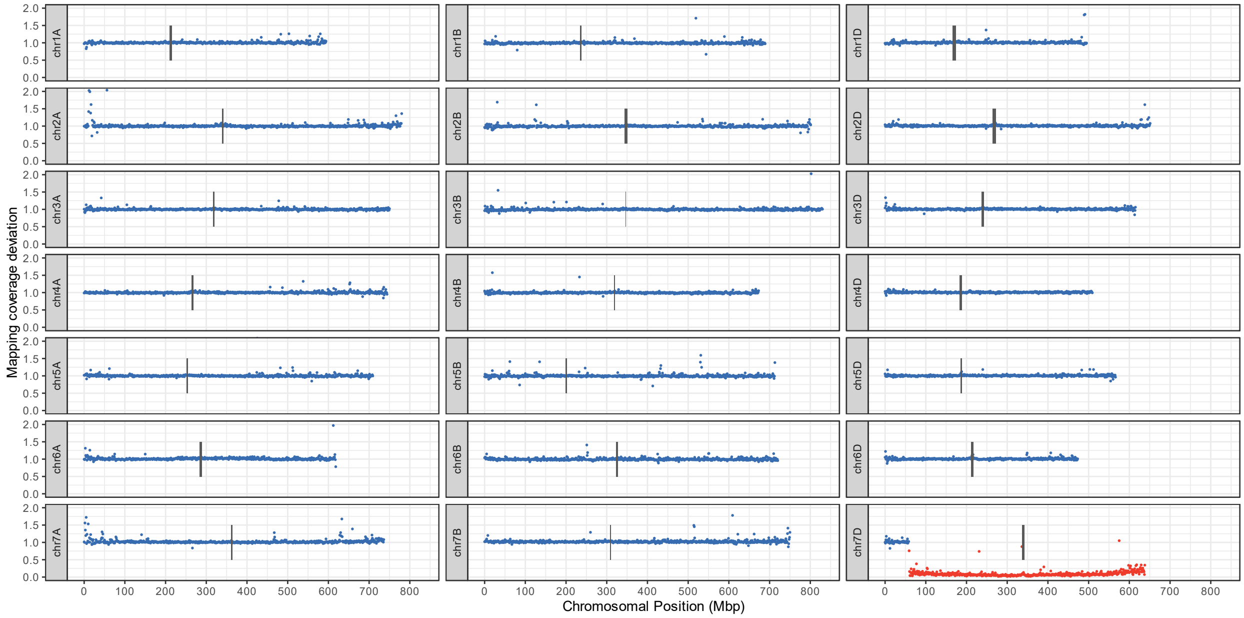
Mapping coverage deviation
Chromosomal Position (Mbp)
chr1A
chr1B
chr1D
chr2A
chr2B
chr2D
chr3A
chr3B
chr3D
chr4A
chr4B
chr4D
chr5A
chr5B
chr5D
chr6A
chr6B
chr6D
chr7A
chr7B
chr7D

DH124

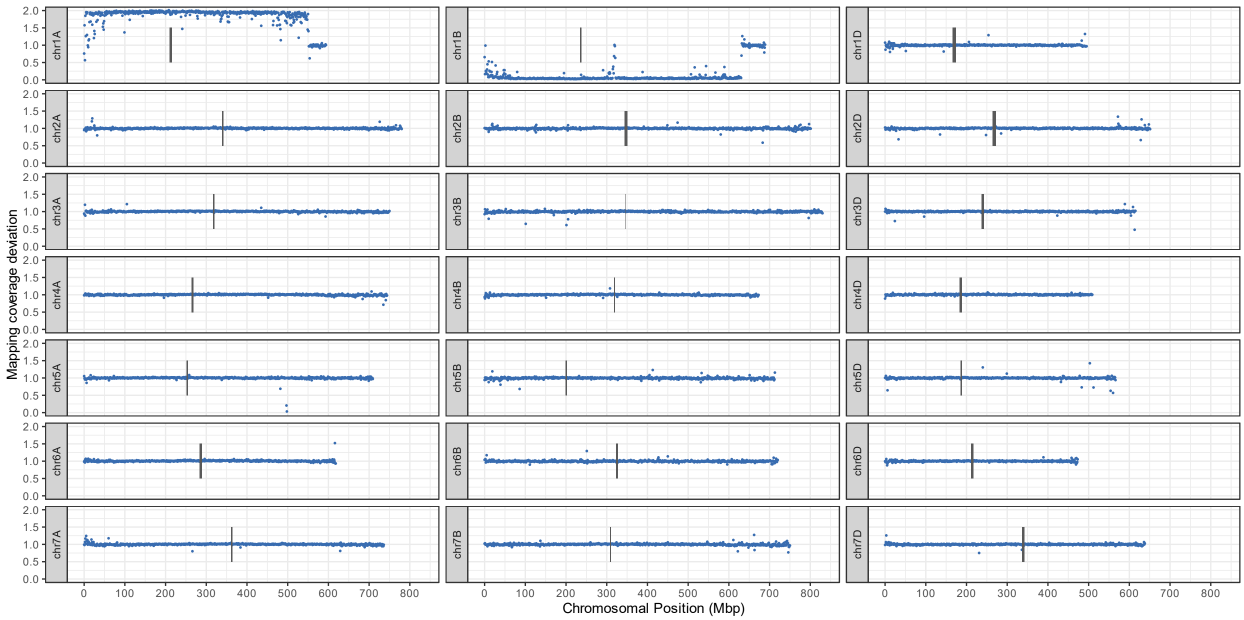
Mapping coverage deviation
Chromosomal Position (Mbp)
chr1A
chr1B
chr1D
chr2A
chr2B
chr2D
chr3A
chr3B
chr3D
chr4A
chr4B
chr4D
chr5A
chr5B
chr5D
chr6A
chr6B
chr6D
chr7A
chr7B
chr7D

DH161

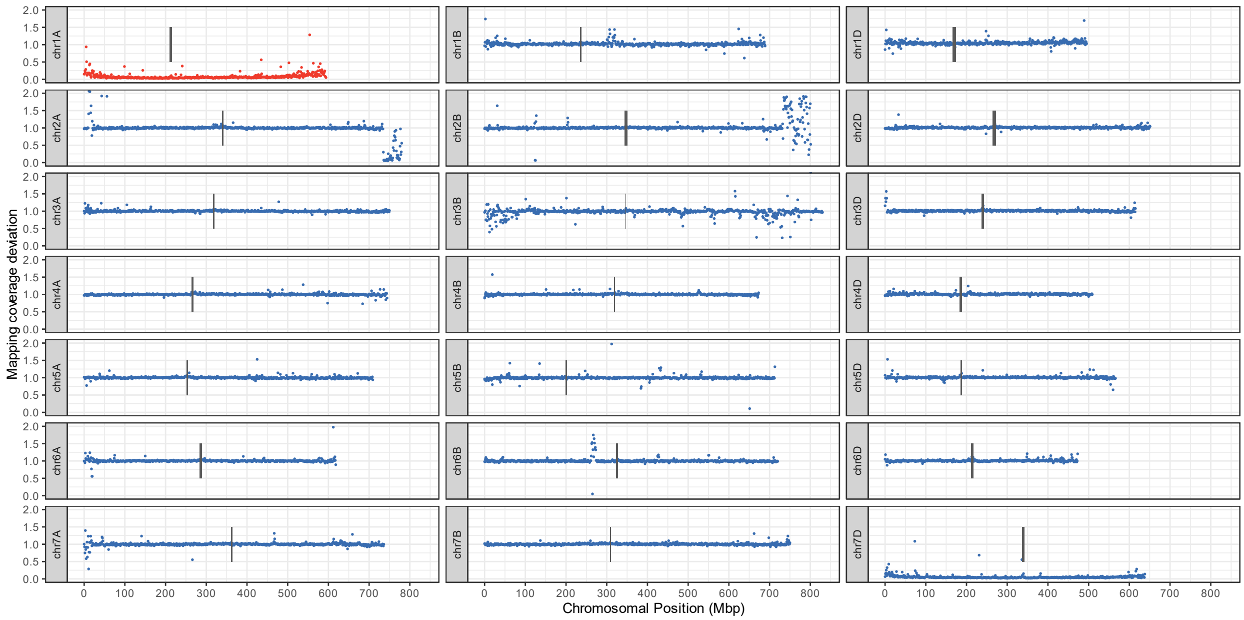
Mapping coverage deviation
Chromosomal Position (Mbp)
chr1A
chr1B
chr1D
chr2A
chr2B
chr2D
chr3A
chr3B
chr3D
chr4A
chr4B
chr4D
chr5A
chr5B
chr5D
chr6A
chr6B
chr6D
chr7A
chr7B
chr7D

DH355

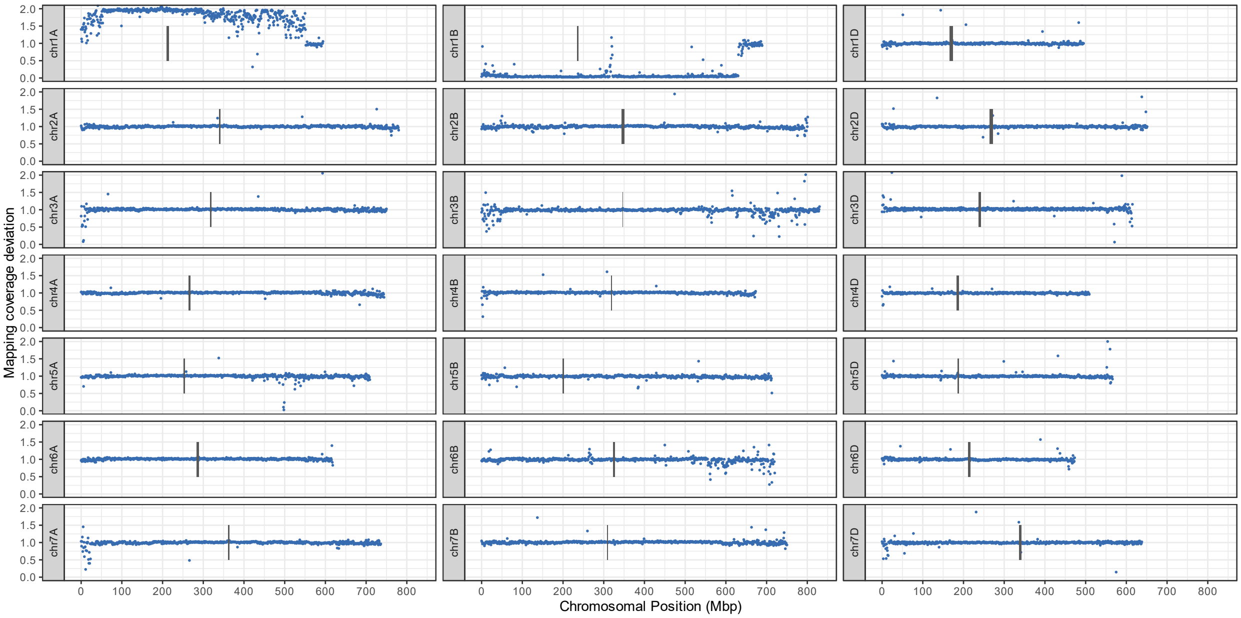


BC2F420

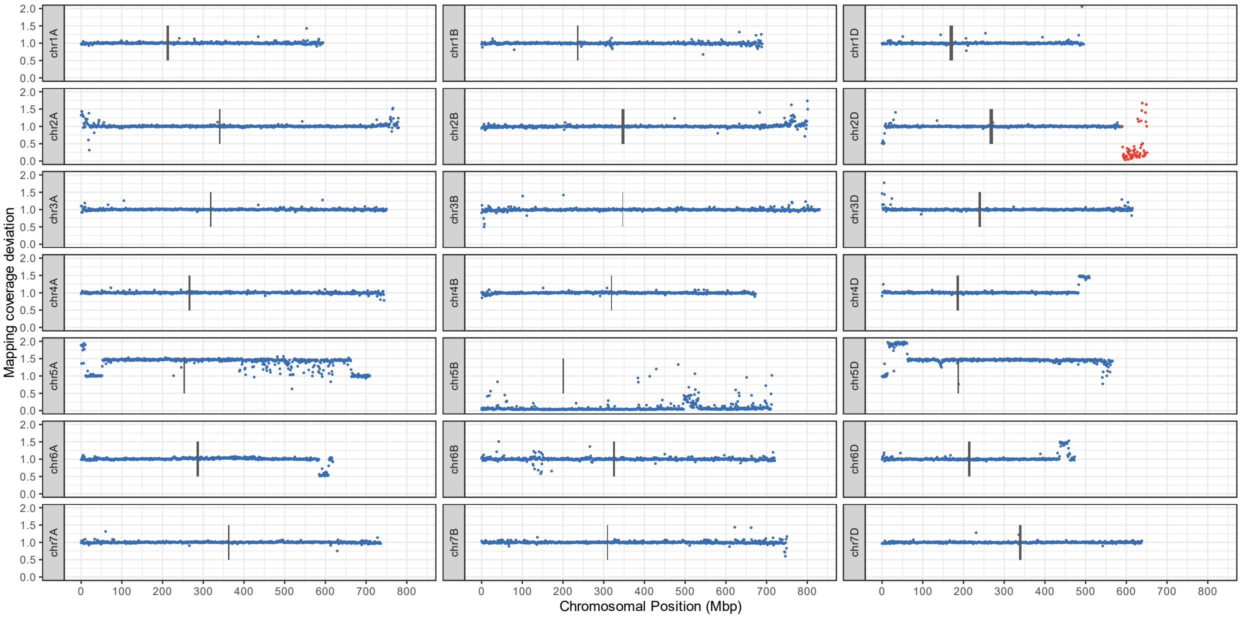

BC3F326

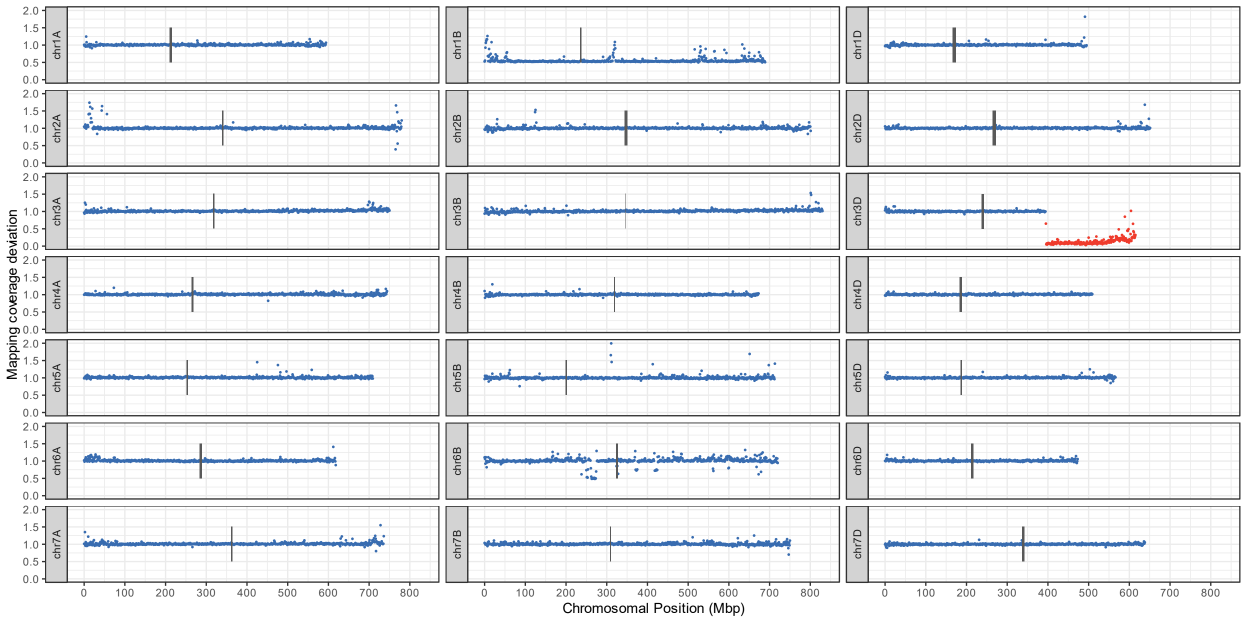


DH195

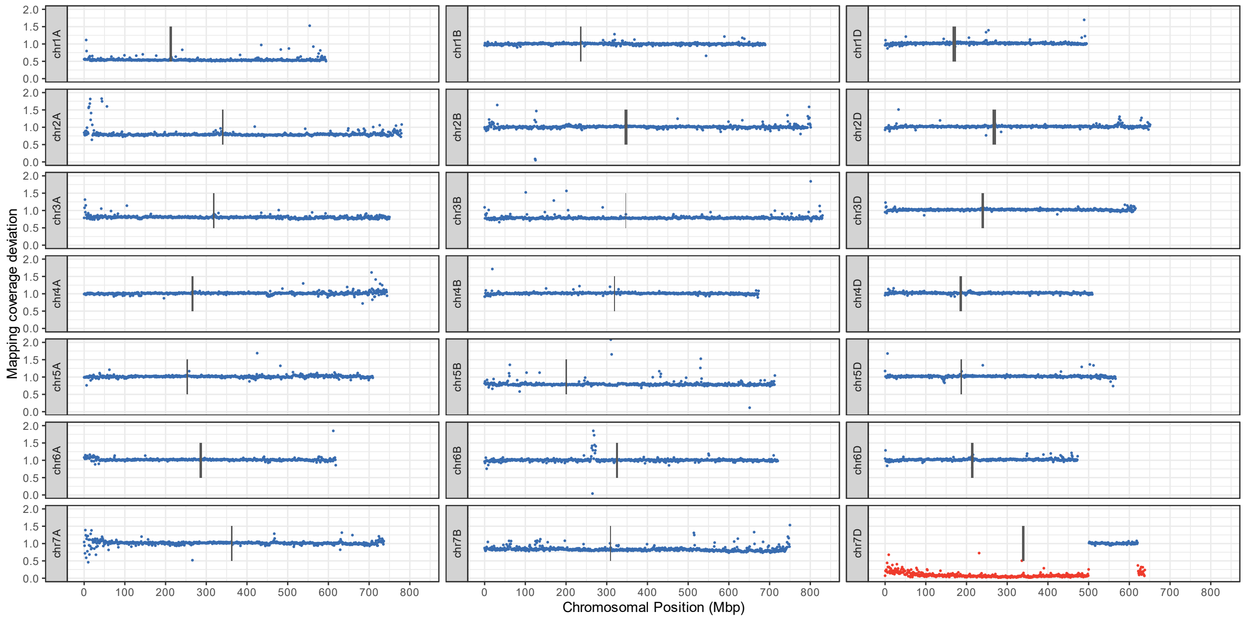

DH202

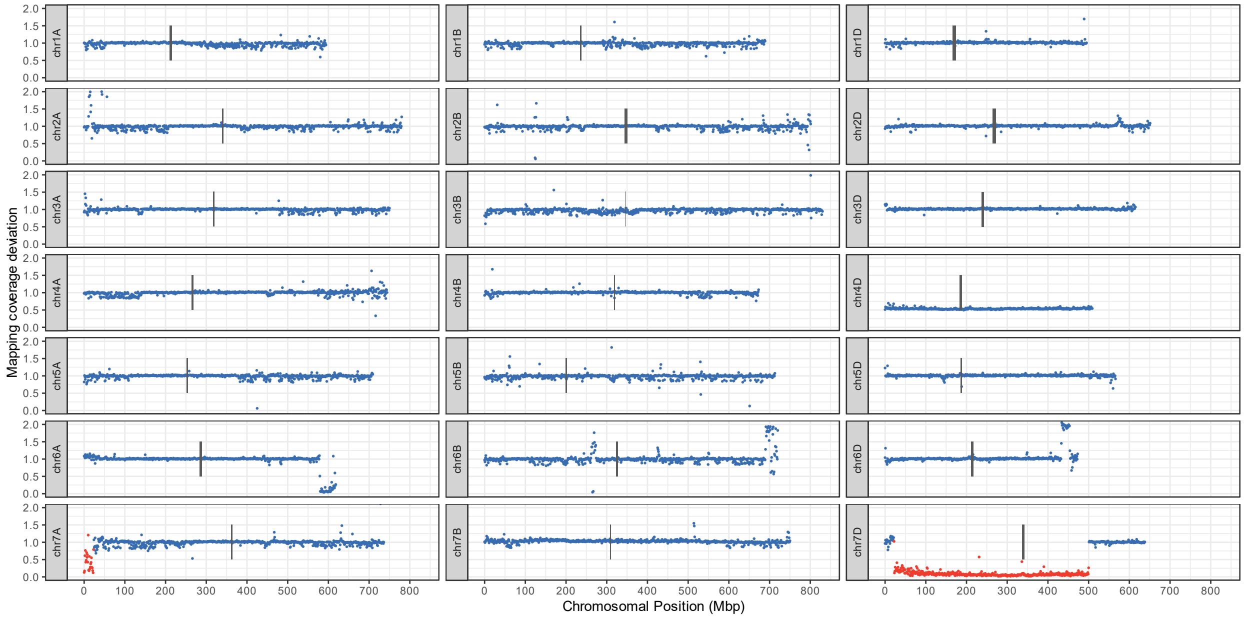


BC3F45

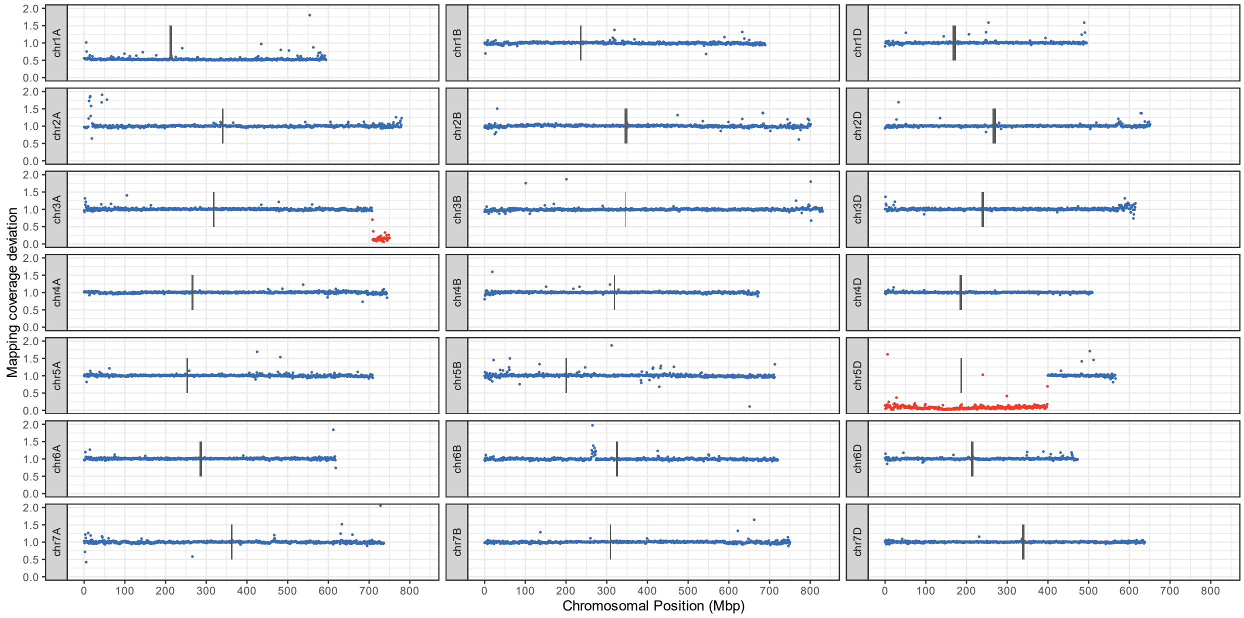

BC3F46

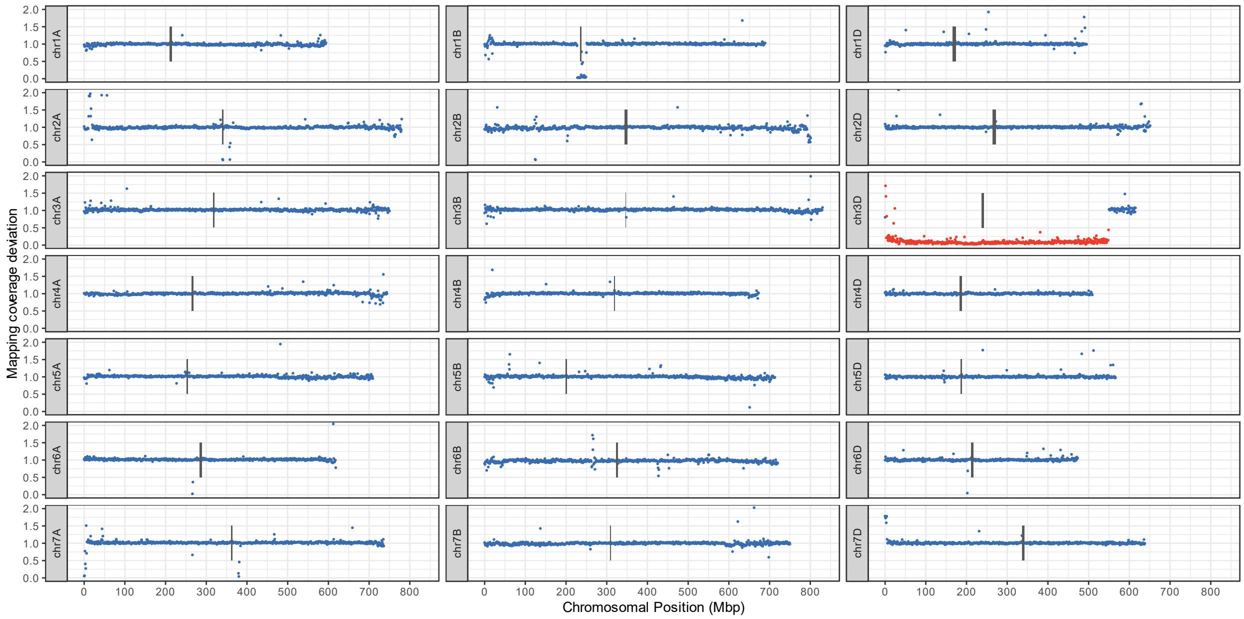


Fig. S1 – Whole genome macro-level plot for all 17 hexaploid wheat/*Am. muticum* introgression lines. Each point represents the deviation in mapping coverage compared to the parent lines in 1Mbp windows across Chinese Spring RefSeq v1.0. Windows within assigned *Am. muticum* introgression blocks are coloured red.

Fig. S2:

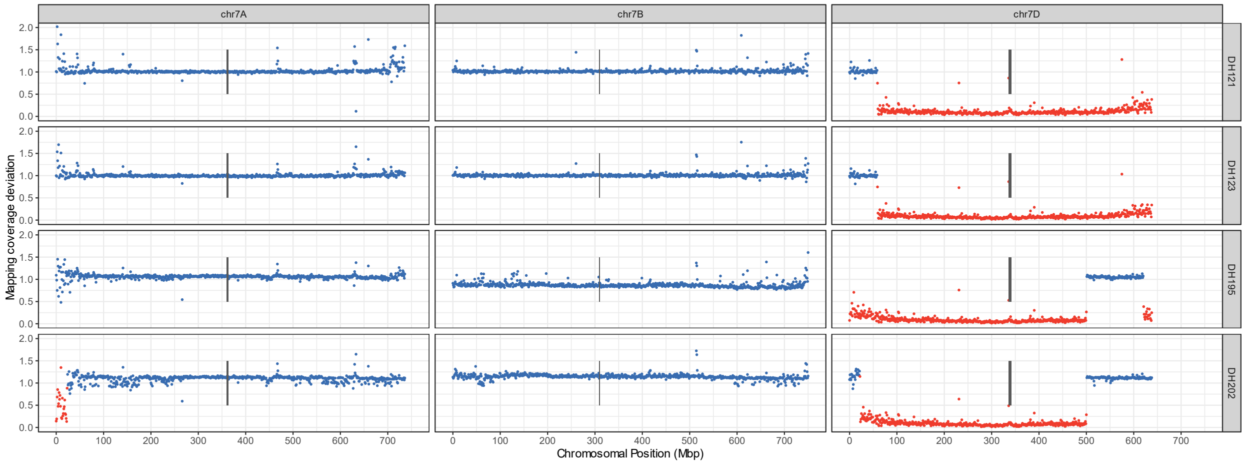


Figure S2: Macro structure of chr7D in four introgression lines (two DH pairs: DH195+DH202, and DH121+DH123). Each point represents the deviation in mapping coverage compared to the parent lines in 1Mbp windows across Chinese Spring RefSeq v1.0. Windows within assigned *Am. muticum* introgression blocks are coloured red. Centromere regions indicated by vertical black bars. These lines provide a good example of how this approach can position segments and resolve overlaps between lines. DH202 appears to have undergone a chr7D-chr7A translocation during crossing which causes the homozygous signal of introgression on chr7A rather than on chr7D. This arises from reads from the chr7D introgression aberrantly mapping to chr7A that can produce a homozygous signal due to the absence of wheat DNA at start of chr7A. Equally, the presence of chr7D wheat DNA on chr7A prevents homozygous *Am. muticum* SNPs, resulting in this region getting filtered out. We can be confident that the large segment in DH202 is unbroken as it appears in DH195 from GISH information and from the deletion at the start of chr7A and retention of the homoeologous region on chr7D.

Fig. S3.

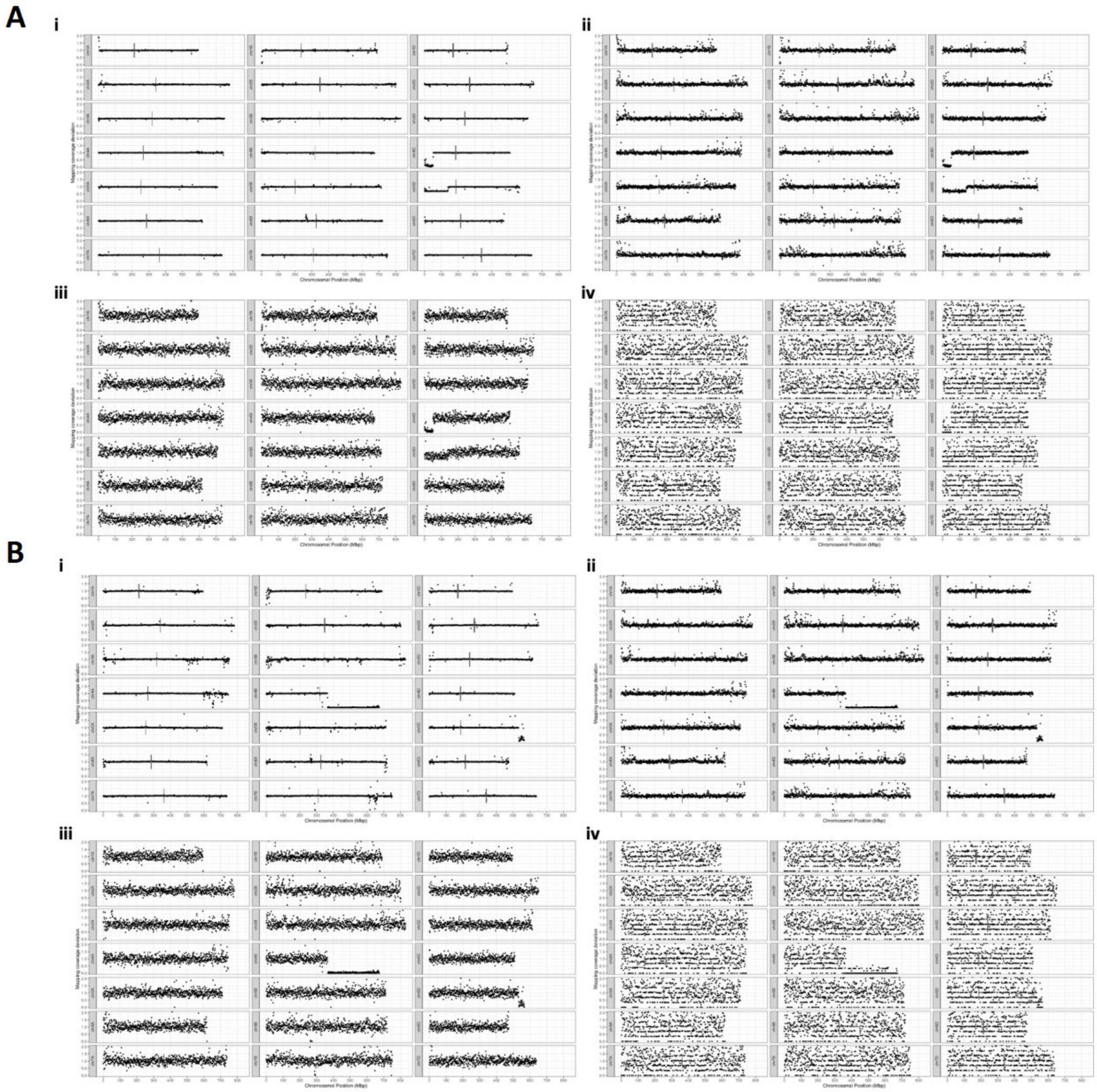


Fig. S3. Minimum required sequencing depth to uncover introgressed segments in introgression lines. Each point represents the deviation in mapping coverage compared to the parent lines in 1Mbp windows across Chinese Spring RefSeq v1.0 for: **A** DH65 and **B** DH92. Downsampled to i 1x; **ii** 0.1x; **iii** 0.01x; **iv** 0.001x.

Fig. S4

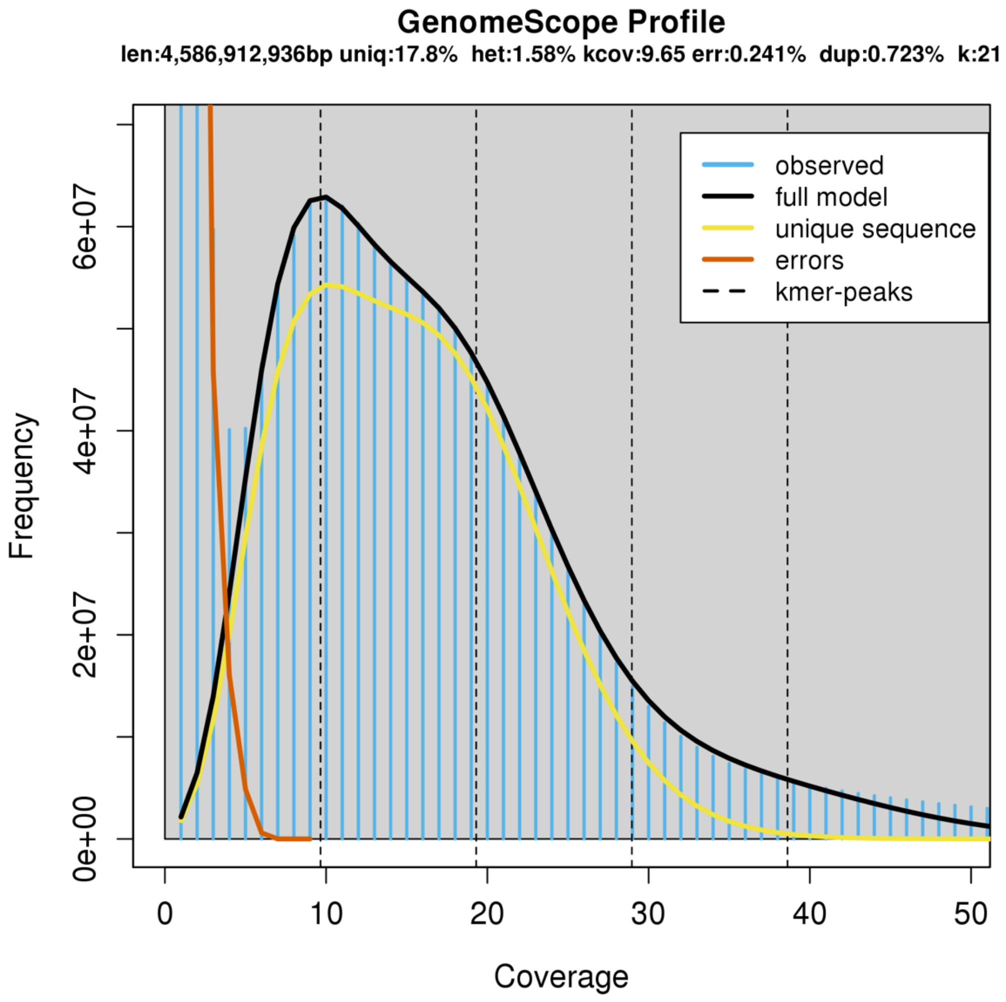


Fig. S4: k-mer distribution of *Am. muticum* Illumina paired-end short reads used to estimate genome size. K-mers counted using Jellyfish with k=21 and processed using GenomeScope 2.0, with parameters kmer length=21, read length=250, max k-mer coverage=10000000.

Fig. S5

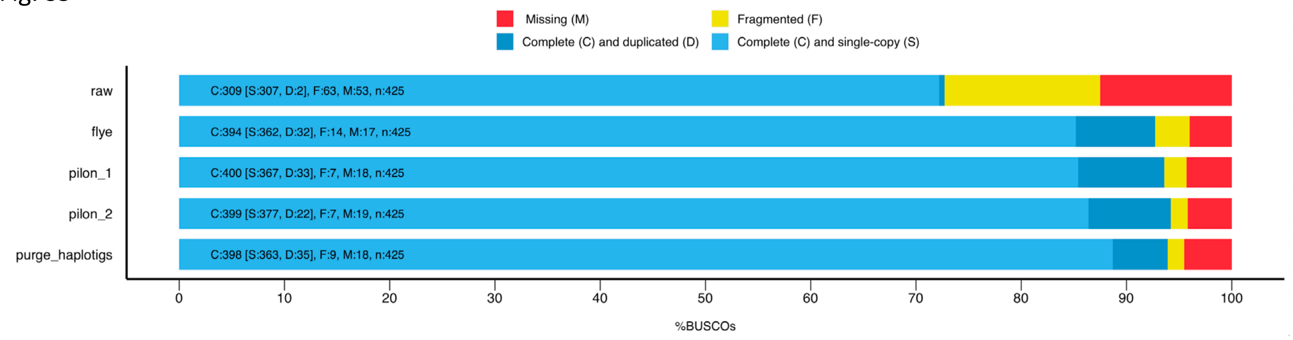


Fig. S5: BUSCO results using the viridiplantae_odb10 dataset after each sequential round of the assembly. Raw = unpolished output from flye, flye = polishing with Oxford Nanopore long reads using flye's built-in tool; pilon_1 = one round of polishing using Illumina paired-end short reads; pilon_2 = two rounds of polishing using Illumina paired-end short reads; purge_haplotigs = removal of uncollapsed haplotigs.

Fig. S6

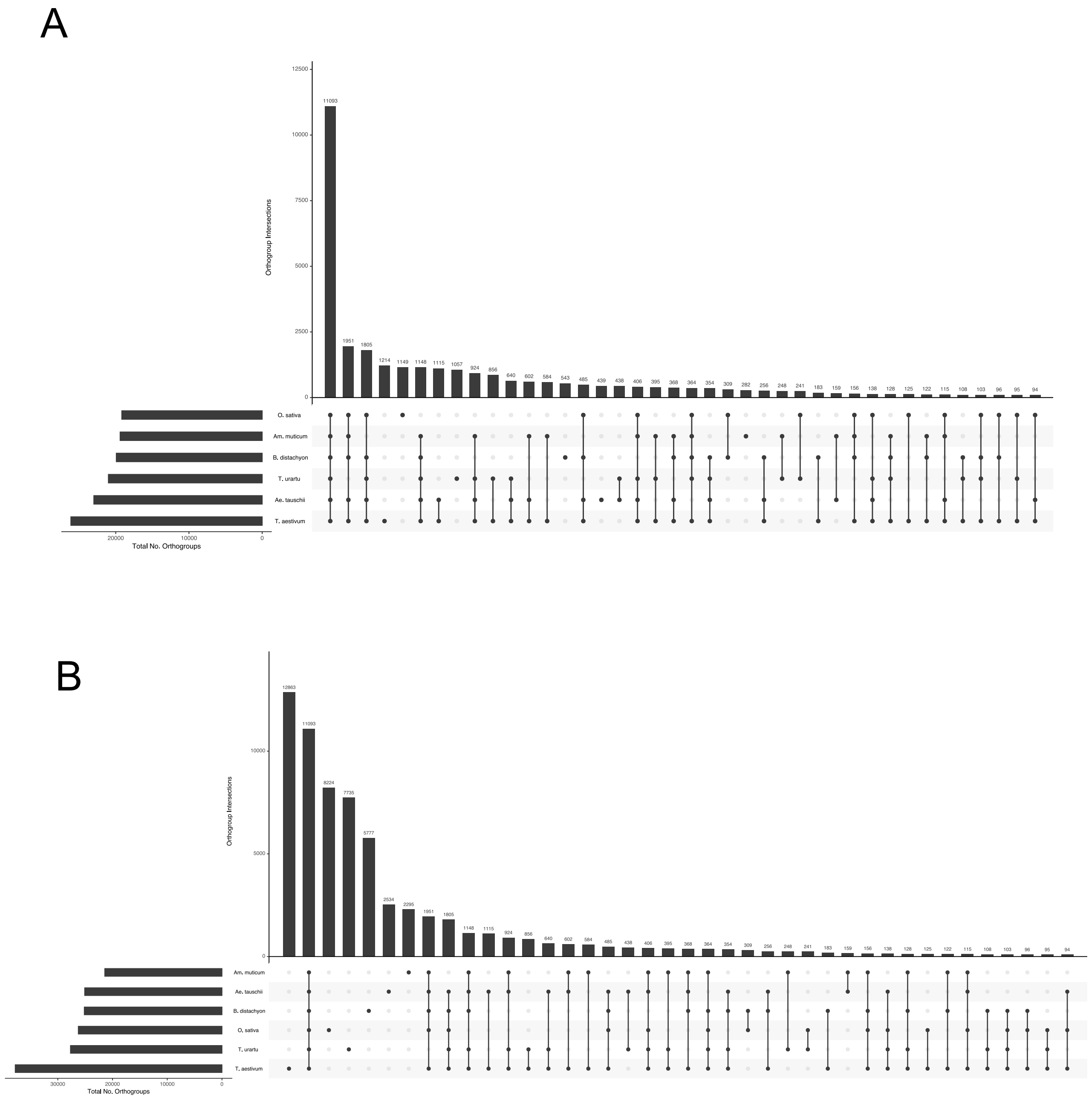


Fig. S6: Interspecies intersection of orthogroups produced by OrthoFinder. **A** Only considering orthogroups with 2 or more members **B** unassigned genes included as single-gene orthogroups

Fig. S7

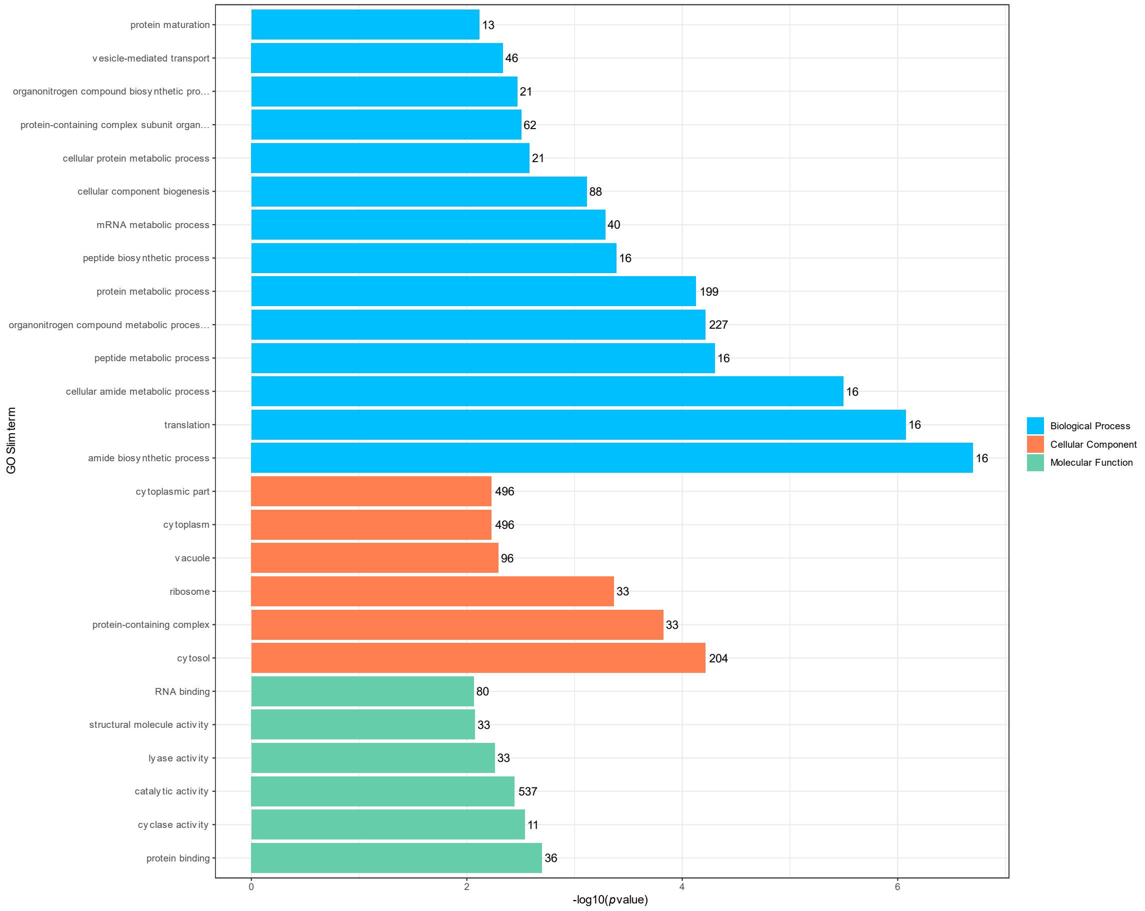


Fig. S7: GO Slim terms enriched in novel *Am. muticum* genes.

Fig. S8.

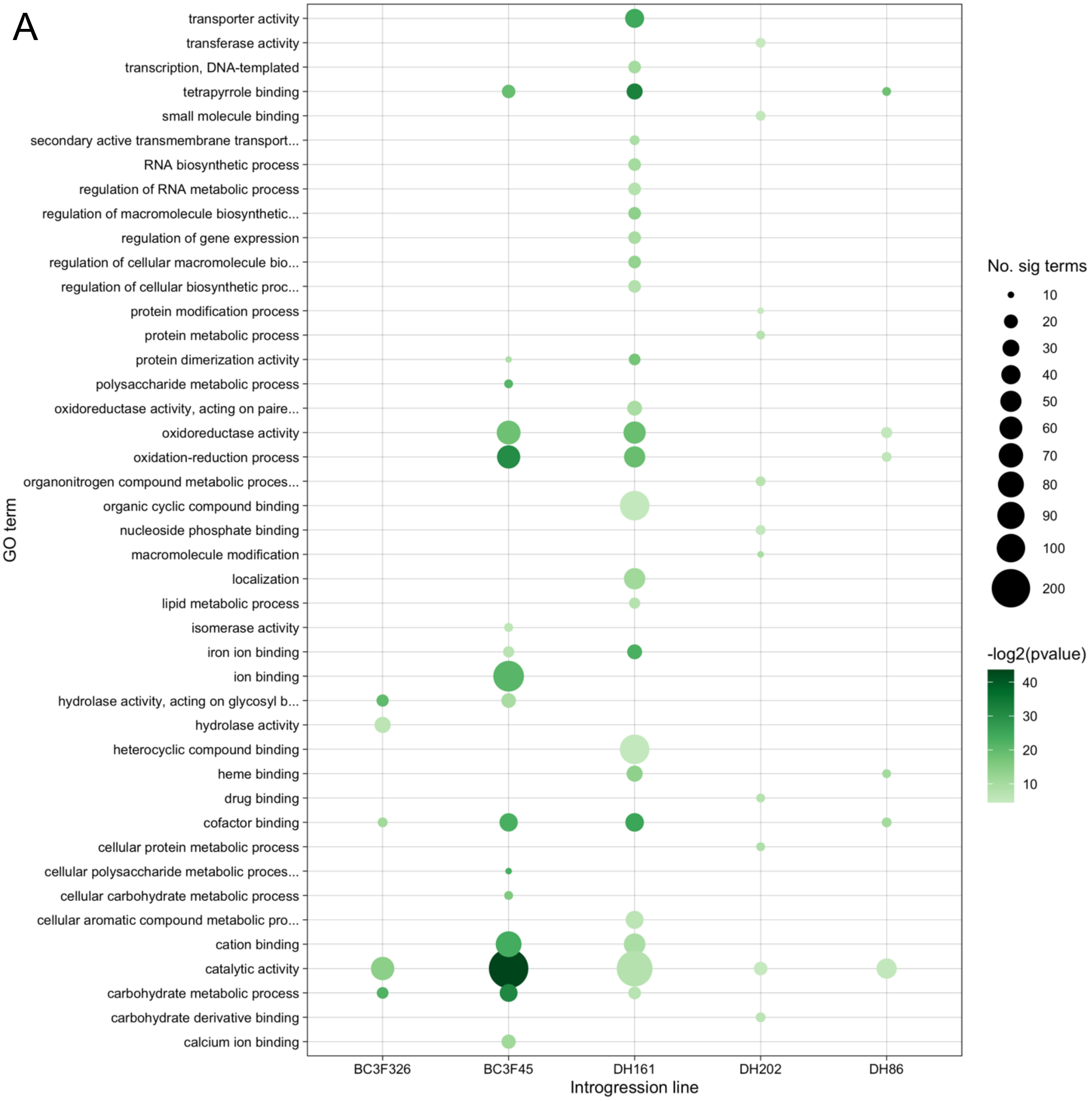

A
GO term
transporter activity
transferase activity
transcription, DNA-templated
tetrapyrrole binding
small molecule binding
secondary active transmembrane transport...
RNA biosynthetic process
regulation of RNA metabolic process
regulation of macromolecule biosynthetic...
regulation of gene expression
regulation of cellular macromolecule bio...
regulation of cellular biosynthetic proc...
protein modification process
protein metabolic process
protein dimerization activity
polysaccharide metabolic process
oxidoreductase activity, acting on paire...
oxidoreductase activity
oxidation-reduction process
organonitrogen compound metabolic proces...
organic cyclic compound binding
nucleoside phosphate binding
macromolecule modification
localization
lipid metabolic process
isomerase activity
iron ion binding
ion binding
hydrolase activity, acting on glycosyl b...
hydrolase activity
heterocyclic compound binding
heme binding
drug binding
cofactor binding
cellular protein metabolic process
cellular polysaccharide metabolic proces...
cellular carbohydrate metabolic process
cellular aromatic compound metabolic pro...
cation binding
catalytic activity
carbohydrate metabolic process
carbohydrate derivative binding
calcium ion binding
BC3F326
BC3F45
DH161
DH202
DH86
Introgression line
No. sig terms
10
20
30
40
50
60
70
80
90
100
200
-log2(pvalue)
40
30
20
10

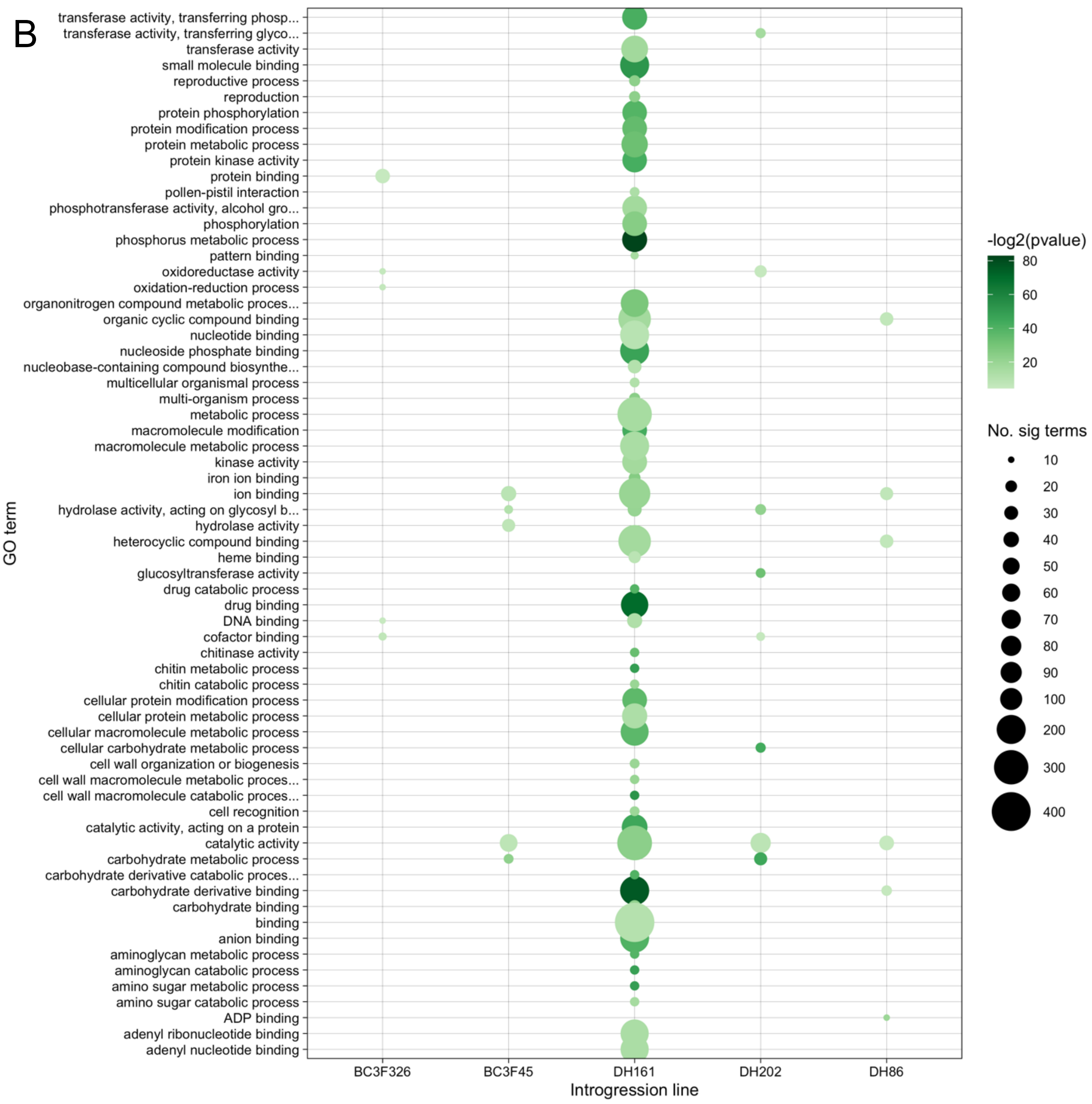


Fig. S8: GO terms enriched in differentially expressed wheat genes. GO term enrichment was performed for genes that were called as differentially expressed and within regions with coverage deviation between 0.8 and 1.0 and not called as introgressed. Enriched terms were included on bubble plot if number of genes with that term >= 10. This was performed separately for genes that were **A** downregulated and **B** upregulated.

Fig. S9.

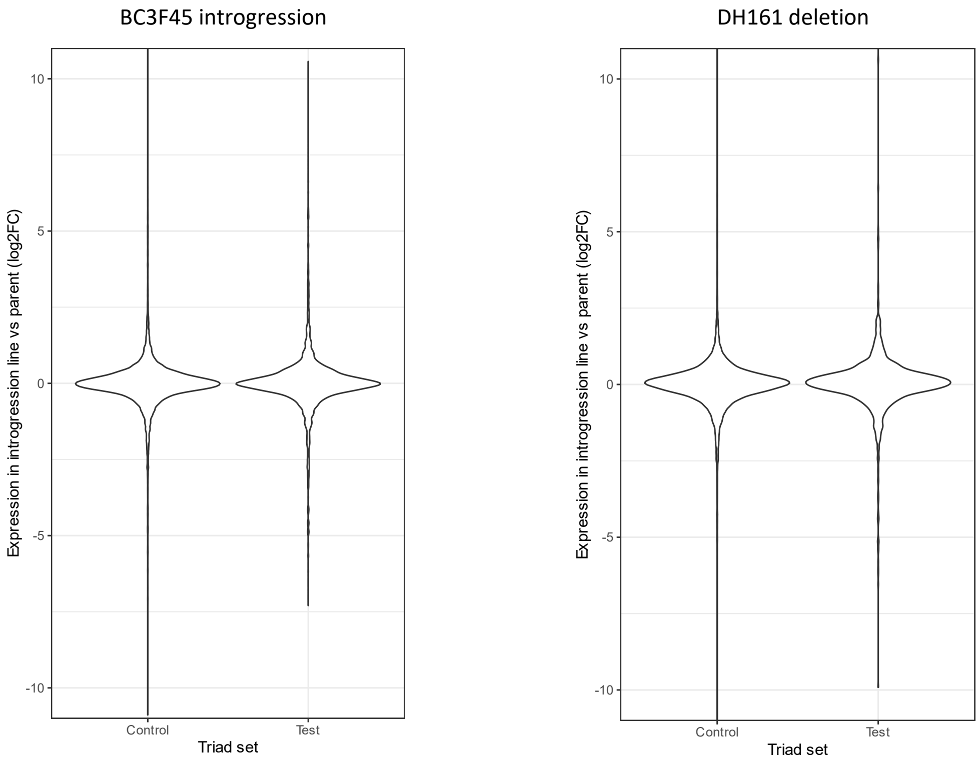


Fig. S9: Testing compensation in homoeologue expression following deletion or introgression. Distribution of log2FC (introgression line vs parent) of A and B homoeologues of triads where the A and B homoeologues are within normal coverage regions, either where the D homoeologue has been replaced with an introgressed *Am. muticum* gene (test) or where the D homoeologue is within a normal coverage region and not called as differentially expressed (control)
