## Supplementary methods for "Whole genome sequencing uncovers the structural and transcriptomic landscape of hexaploid wheat/*Ambylopyrum muticum* introgression lines"

**Methods S1: Introgression line production:**

As described in (King et al., 2019, 2017), *Am. muticum* accessions 2130004/2130012 were crossed with Pavon76 or Chinese Spring. The F1 interspecific hybrid was backcrossed 3 times, once with Paragon, Pavon76 or Chinese Spring and twice with Paragon. To make homozygous lines in which introgressions are stably inherited, the resulting BC3 lines were either pollinated with maize and treated with colchicine to generate doubled haploids (DH) (King et al., 2019) or were selfed 2 or 3 times.

**Method S2. DNA extraction and whole-genome sequencing:**

*Am. muticum* introgression lines and wheat parents were grown in a growth room (16h, 21°C day/8h, 18°C night). Genomic DNA from young leaves was isolated using extraction buffer (0.1 m Tris–HCl pH 7.5, 0.05 m EDTA pH 8.0, 1.25% SDS). Samples were incubated at 65 °C for 1 h before being placed on ice and mixed with ice-cold 6 m NH4C2H3O for 15 min. The samples were then spun down, the supernatant was mixed with isopropanol to pellet the DNA and the isolated DNA was treated with RNase A then purified with phenol/chloroform. DNAs were dissolved in TE (10mM Tris-HCl pH8.0, 0.1mM EDTA). PCR-free libraries were produced from this DNA with >600bp insert sizes (gel size-selection). These were sequenced on Illumina NovaSeq 6000 S4 flowcells to produce 150bp paired-end reads.

**Method S3. Read mapping and SNP calling:**

Reads were mapped to the Chinese Spring reference genome RefSeq v1.0 (International Wheat Genome Sequencing Consortium (IWGSC) et al., 2018) using BWA-MEM (Li, 2013) with the -M parameter to enable duplicates to be marked. The alignment was filtered using samtools (Li et al., 2009): supplementary alignments, improperly paired reads, and non-uniquely mapped reads (q <= 10) were removed. PCR duplicates were detected and removed using Picard’s MarkDuplicates (DePristo et al., 2011). Variants were called using samtools mpileup (Li et al., 2009) and bcftools call (Li, 2011) using the multiallelic calling model -m. INDELs were removed and SNPs retained with a quality score >= 30 and a read depth >= 10 for the parental lines and >= 5 for the introgression lines (mean sequencing depth of introgression lines). Introgression line SNPs were further filtered as follows: homozygous SNPs were kept if 5 or more reads supporting the alternative allele and the allele frequency (AF) was 1; heterozygous SNPs were kept if 3 or more reads supporting each allele. Sites with 3 or more alleles were removed.

**Method S4. Producing *Am. muticum*-specific SNPs**

*Am. muticum* SNPs were retained as genome-specific if not shared with Paragon or Pavon76. At heterozygous sites where one allele was specific to *Am. muticum*, the specific allele was retained. If two alternative alleles were present, both were kept if both were specific to *Am. muticum*.

**Method S5. Assigning coverage deviation blocks as *Am. muticum*:**

1Mbp windows containing >= 55 homozygous *Am. muticum* specific SNPs and a ratio of homozygous to heterozygous *Am. muticum* specific SNPs >= 4 that fell within a coverage deviation block were classified as candidate *Am. muticum* windows. Coverage deviation blocks with >= 14% windows assigned as *Am. muticum* using the parameters above were classed as an introgressed segment. These parameters are adjustable and were chosen because they revealed previously known introgression segments and didn’t falsely class any known deletions as introgressions. The coverage deviation values in the 100Kbp windows up and downstream of either end of the introgression blocks were examined to determine where the coverage deviation starts to decrease. The first 100Kbp window either side that falls below coverage deviation of 0.7 was assigned as the border window of the segment for manual inspection.

**Method S6. KASP genotyping:**

Genomic DNA was isolated from leaf tissue of 10-day old seedlings in a 96-well plate as described by Thomson and Henry (Thomson and Henry, 1995). All DH lines were genotyped alongside the three parental wheat genotypes (Chinese Spring, Paragon and Pavon 76) and the two *Am. muticum* accessions as controls. For each KASP™ assay, two allele-specific primers and one common primer were used (Table S4). The genotyping procedure was as described in (Grewal et al., 2020). In summary, the genotyping reactions were set up using the automated PIPETMAX^®^ 268 (Gilson, UK) and performed in a ProFlex PCR system (Applied Biosystems by Life Technology) in a final volume of 5 μl with 1 ng genomic DNA, 2.5 μl KASP reaction mix (ROX), 0.068 μl primer mix and 2.43 μl nuclease free water. PCR conditions were set as 15 min at 94°C; 10 touchdown cycles of 10 s at 94°C, 1 min at 65–57°C (dropping 0.8°C per cycle); and 35 cycles of 15 s at 94°C, 1 min at 57°C. Fluorescence detection of the reactions was performed using a QuantStudio 5 (Applied Biosystems) and the data analysed using the QuantStudio^TM^ Design and Analysis Software V1.5.0 (Applied Biosystems).

**Method S7. Estimating genome size:**

Filtered nanopore reads were mapped back to the polished genome assembly using minimap2 (Li, 2018) with parameters -x asm5 and filtered to retain uniquely mapping reads with mapQ >= 10. We computed mean coverage across single-copy BUSCO genes found in the assembly. The estimated genome size was calculated as the total number mapped bases / mean coverage across single-copy genes, using post-filtering numbers. k-mers were counted in the raw Illumina paired-end reads using Jellyfish (Marçais and Kingsford, 2011) with k=21. These k-mers were analysed using GenomeScope 2.0 (Ranallo-Benavidez et al., 2020) with parameters kmer length=21, read length=250, max kmer coverage=10000000 (Fig. S4).

**Method S8. Repeat annotation and masking:**

A de novo library of transposable elements was produced using EDTA v1.9.5 (Ou et al., 2019), using cds sequences from *T. aestivum* to exclude protein-coding regions from the library. The resulting library was aligned to the assembly using blastn (Camacho et al., 2009) and sequences with fewer than 3 full length hits (defined as over 80.0% similarity across more than 80.0% of the length of the element) were removed and remaining sequences were clustered using cd-hit-est v4.6.7 (Li and Godzik, 2006) with parameters aL 0.8 aS 0.8 -c 0.8 to remove redundant sequences. The resulting library was used to mask the genome assembly using RepeatMasker v4.07 (Chen, 2004) with parameters -s -no_is -norna -nolow -div 40 -cutoff 225.

**Method S9. Gene annotation:**

*Ab initio*, protein homology and transcriptome evidence were combined to predict protein-coding genes in the *Am. muticum* assembly. First, 4 biological replicates of *Am. muticum* root and shoot mRNA extracted and sequenced as in method Sx. and were trimmed using Trimmomatic (Bolger et al., 2014) with the parameters ILLUMINACLIP:BBDUK_adaptor.fa:2:30:12 SLIDINGWINDOW:4:20 MINLEN:20 AVGQUAL:20. The trimmed RNA reads were mapped to the masked genome using STAR (Dobin et al., 2013). Transcripts were assembled using four independent reference-guided approaches; Trinity (Grabherr et al., 2011); StringTie (Pertea et al., 2015), cufflinks (Trapnell et al., 2012) and CLASS2 (Song et al., 2016). Transdecoder (Haas et al., 2013) was used on each set of transcripts to produce coding ORFs. PORTCULLIS (Mapleson et al., 2018) was used to produce splice information. Uniprot (UniProt Consortium, 2019) reference proteomes for *T. aestivum*, *Ae. tauschii*, *T. turgidum*, *Oryza sativa*, *B. distachyon*, and *A. thaliana* were aligned to the genome using blastx (Camacho et al., 2009). Mikado (Venturini et al., 2018) was used to merge and refine the transcripts produced by each tool, aided by the splice site information and the protein homology evidence. The same uniprot reference proteomes as above were aligned to the masked genome using TBLASTN (Camacho et al., 2009). Hits with an e-value below 1e-5 and sequence identity greater than 75% within 20Kb were merged using bedtools merge (Quinlan and Hall, 2010). These regions were searched using exonerate (Slater and Birney, 2005) to refine protein alignments. To produce a set of transcript annotations to train Augustus (Hoff and Stanke, 2019), Mikado protein coding genes were filtered to retain multi-exon genes with < 80.0% amino acid identity with any other protein, not have a stop codon in the ORF, and be at least 500bp away from another gene. Of the remaining transcripts, the 2000 with the highest mikado score were retained and randomly split into 1800 training genes and 200 testing genes. The AUGUSTUS HMM was trained using these transcripts using etraining followed by 10-fold cross validation using optimize_augustus.pl. To produce initial *ab initio* gene predictions, Augustus was run using this HMM, along with intron hints produced from the STAR bam file using bam2hints from Augustus. EvidenceModeler (Haas et al., 2008) was used to produce a set of HC gene models with *ab initio*, transcriptome, and protein homology evidence. This set of HC genes was used to retrain Augustus as before, as well as GlimmerHMM (Majoros et al., 2004). Augustus, SNAP (Korf, 2004) (using the rice HMM) and GlimmerHMM were run to produce final ab initio gene predictions. These predictions were incorporated with the transcript and protein homology evidence using EvidenceModeler to produce final gene models with the following weightings: mikado transcriptome, weight 14; Augustus gene models - weight 4; exonerate protein evidence - weight 7; SNAP gene models - weight 1; GlimmerHMM gene models - weight 2. Predicted proteins blasted to the TREP database of transposable elements and to TREMBL proteins. Genes were classed as high-confidence if they had a complete protein-coding gene model, transcript evidence from one or more of the transcriptome assembly methods, and their encoded protein had one or more significant hits to TREMBL (Bairoch and Apweiler, 1997) and no significant hit to TREP (Wicker et al., 2002). Gene functions were assigned using eggnog 5.0 (Huerta-Cepas et al., 2019).

**Method S10. mRNA extraction and sequencing:**

Seeds were germinated and grown to the 4^th^ leaf stage in a growth chamber set to 22C, 16-hour days. Plants were well watered and supplemented with nutrients. The 4^th^ leaf was extracted and snap frozen in liquid nitrogen within 20 seconds of sampling and stored at -80C. Three reps of each line were sampled individually. Each sample was ground in a pestle and mortar using liquid nitrogen and a sub-set taken for Total RNA extraction using Qiagen RNeasy plant mini kit (Qiagen, Germany). DNA was removed using a digestion step with RNase-free DNase (Qiagen, Germany). Ratios of 260:280 were measured using a Nanodrop (ThermoFisher Scientific) and samples were only used with ratios between 1-9 and 2.1. Only samples with RNA integrity number (RIN) >8 and a concentration of 50-200ng/ul (Agilent Bioanalyzer) were sequenced. Library preparation was conducted using automated NEBNext Ultra II Directional RNA-Seq library construction with Poly-A selection and was sequenced in triplicate on an Illumina NovaSeq 6000 S2 flowcell by Genomic Pipelines at the Earlham Institute to produce 150bp paired-end reads

**Method S11. Identifying resistance genes:**

*Am. muticum* NLR genes were identified using two parallel methods. The first was to scan high and low confidence gene models for Pfam domains found in complete NLR genes using hmmscan from the HMMER package (Finn et al., 2011). The second was to scan the entire genome de novo for complete, functional NLR protein-coding regions using NLRAnnotator (Steuernagel et al., 2020). This allowed several additional NLR genes whose gene model had been filtered out or was never constructed to be recovered. Other types of genes that have been previously implicated in stripe rust resistance (Zheng et al., 2020) (ABC transporters, Pkinases, hexose transporters, wheat Kinase-START genes, tandem kinase-pseudokinase proteins) were identified from the eggnog annotation and validated manually using hmmscan and pfam domains.
